# Protein language models learn the underlying accessibility of mutations alongside protein fitness landscapes

**DOI:** 10.64898/2026.07.06.736753

**Authors:** Oscar A MacLean, Kieran Lamb, Spyros Lytras, Laura Mojsiejczuk, Ke Yuan, Joseph Hughes, David L Robertson

**Affiliations:** MRC-University of Glasgow Centre for Virus Research, School of Infection and Immunity, Glasgow, UK; Antigen Evolution and Design Lab, Department of Structural Biology and Chemistry, Institut Pasteur, Paris, France; School of Cancer Sciences, University of Glasgow, Glasgow, UK; Cancer Research UK Scotland Institute, Glasgow, UK

## Abstract

Protein language models (PLMs) score the effects of amino acid replacements as pseudo-probabilities, which are widely utilised to map protein fitness landscapes. However, because their training data relies on natural amino acid sequences, these models conflate protein structural constraints with nucleotide mutation biases and codon accessibility. Using the rapid emergence of the divergent influenza A H3N2 K lineage as a stress test, we investigate how base PLMs (ESM-2 and ESM-C) versus fine-tuned versions of these models capture mutational processes. We systematically implement a parameter sweep to explicitly couple (or decouple) empirical nucleotide mutational supply from PLM-assessed amino acid substitution pseudo-probabilities across evolutionary forecasting tasks. We find that base PLMs implicitly learn generic nucleotide-level mutational constraints, an effect strongly amplified by virus-specific fine-tuning. Incorporating explicit mutational accessibility significantly improves the binary prediction of which amino acid replacements will be observed. Conversely, when predicting the observed frequency of already circulating variants that have emerged, adding mutational supply degrades performance, confirming that selection dominates post-emergence dynamics. Our results demonstrate that current PLM pseudo-probabilities are a composite metric that conflates protein structural fitness with historical biases in mutational supply. Fine tuning the models on closely related viral sequences greatly strengthens the models’ mutational accessibility knowledge, even learning which synonymous codons are being used at which site. Correlations of these fine-tuned models to deep mutational scanning data are improved by downweighting replacements by a function of their mutational accessibility. This supports explicitly decoupling these independent evolutionary processes to optimise predictive accuracy for real-world pathogen forecasting and help isolate protein fitness for synthetic design pipelines.

## Introduction

Protein language models (PLMs) provide a powerful new tool to quantify the structural and evolutionary constraints of proteins (Bepler and Berger 2021; Elnaggar et al., 2022; Lin et al., 2023). The major output from these models is pseudo-probabilities for all possible sequence amino acid residue changes. These represent the model’s inferred chance of observing a given amino acid token at that site for a given embedded sequence, which can be applied to help understand virus evolution (Hie et al., 2021;Ito et al., 2025a; Lamb et al., 2026; Yang et al., 2026). These are often interpreted as a proxy for the protein’s fitness landscape, reflecting functional constraints imposed by natural selection.

The training data for these models, however, are protein sequences governed by complex evolutionary processes. Evolution is not driven purely by the protein’s functional constraints and associated fitness landscapes; it is strongly influenced by the biased nature of nucleotide mutations (Yampolsky & Stoltzfus, 2001; Stoltzfus, 2006). Certain amino acid replacements are vastly more likely to occur than others. These differences are driven by 1) directional mutation biases, and 2) the fact that certain amino acid replacements require two or three nucleotide mutations rather than one. These two forces constrain the mutational accessibility of amino acid changes for all proteins. Additionally, direct selection at the nucleotide level can also shape accessibility, skew codon usage and distort protein selection inference (De Maio et al., 2021). The lower resulting rate of certain amino acid replacements is directly internalised by PLMs, shaping their learned representations of protein space. Replacements may never be observed in the training data because they have a lethal fitness effect or simply because they are many orders of magnitude less accessible. This accessibility signal confounds the models’ probabilities, and interferes with their utility for both protein structural design and evolutionary prediction purposes. For viral proteins, model pseudo probabilities have been shown to enrich for viable mutations, albeit often to a limited degree (Allman et al., 2025). This limitation is particularly acute in models of antibody evolution from somatic hypermutation. Lightweight models that explicitly account for codon-level mutational accessibility outperform both antibody-specific and general-purpose PLMs in predicting affinity maturation trajectories (Johnson et al., 2025). This mutation bias has been shown to broadly confound PLM pseudo-probabilities across both antibody and general-purpose PLMs, reducing the utility of the models (Matsen et al., 2026). We further explore this in the context of evolutionary forecasting of viral datasets, decomposing the bias across codon space, and investigate how model fine-tuning impacts the bias.

Predicting pathogen evolution remains a critical public health challenge, as demonstrated by influenza. The virus imposes a severe biological and economic toll, causing an estimated 290,000 to 650,000 deaths annually (Iuliano et al., 2018) alongside substantial global economic disruption (Putri et al., 2018). Influenza infections which bypass vaccine immunity are estimated to cost $29 billion in the USA alone (Popovian and Winegarden 2025). The multi-month turnaround required for sampling, testing, and manufacturing makes predicting each year’s dominant influenza strain a persistent challenge, as demonstrated by the sudden emergence of the K lineage during the 2025/2026 season (Kirsebom et al., 2025; Dee et al., 2026).

Recent computational advances have enabled accurate *in silico* prediction of antibody escape from a single influenza Haemagglutinin (HA) sequence (Ito et al., 2025b). This method, trained on ferret data, embeds sequences into 3D antigenic space to predict cross immunity between H3N2 influenza variants. A powerful step alongside this inference would be predicting unobserved sequences with the potential to emerge in the future. However, with 20 possible amino acids at 566 HA sites, the fitness landscape of the HA protein is immense. Relative to the J.2 parental lineage which emerged in late 2023, the more recent K lineage is defined by nine Haemagglutinin mutations (Sabaiduc et al., 2025). The combinatorial nature of mutations creates a vast search space in the fitness landscape at even this level of divergence. Indeed, there are 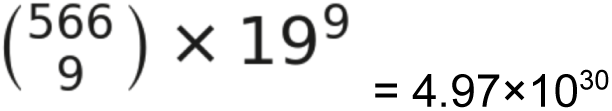 possible sequences with nine mutations from that J.2 node, an unsearchable space, before considering neuraminidase or any of the other encoded proteins.

Predicting evolutionary trajectories requires more than mapping a sequence’s structural fitness; it also requires accounting for the mutational accessibility of the amino acid changes. These probability differences are shaped by the number of nucleotide mutations required, and asymmetric nucleotide mutation biases. Host driven mutational forces impose a strong A->G and T->C mutation bias on influenza genomes (Pauly et al., 2017). Some influenza A amino acid replacements are 6.5×10^11^-fold more likely to occur than others (Pauly et al., 2017; Cano et al., 2022; Gunnarsson and Babu 2023). Consequently, mutational accessibility can dwarf predicted fitness effects of amino acid changes (Gunnarsson and Babu 2023). From a PLM output, the pseudo-probability of the top and bottom amino acid replacements have a probability scale difference of 10^8^. Incorporating this larger 6.5×10^11^ range of mutational accessibility to the PLM predictions of amino acid changes has the potential to reduce the plausible protein space massively, significantly improving forecasting. A concurrent analysis by Efimenko et al. (2026) found a nearly neutral framework using nucleotide mutational spectra explained more than 50% of observed single-nucleotide amino acid substitutions in SARS-CoV-2. Conversely, for predicting the immediate spread of *existing* variation, the mutational accessibility information captured by PLMs could theoretically be a confounding variable, downweighting rare low-accessibility mutations which have already overcome the mutational barrier.

The ability to fine-tune PLMs on certain proteins or protein families has often been shown to improve model performance on prediction tasks (Lafita et al., 2024; Sawhney et al., 2025; Bedford 2026). This process takes a general model’s weights trained on diverse species, and performs additional model training epochs with species specific training data, aiming to switch a generalist model to a specialist. Bedford (2026) found that counting mutations in key regions produced slightly stronger predictions of SARS-CoV-2 fitness flux than PLM embeddings, even in fine-tuned models—though this may be confounded by positive selection saltations producing long branches. A recent paper fine-tuned the ESM-2 protein language model on influenza HA sequences, producing the HA-80 model (80-20 temporal train-test split; 2015 training data cut-off). This model gave higher future predictive power of mutations on downstream phylogenetic nodes (Lytras et al., 2026). However, this additional training with more closely related sequences may exacerbate the mutational bias knowledge implicit within the PLM output. Sequences which are clustered to low identity thresholds will span across longer evolutionary time periods allowing rarer mutational events to accumulate.

Here we aim to disentangle how PLMs learn both protein fitness and the underlying mutation bias, as their training data is inherently confounded by the two forces. The recent emergence of the novel influenza H3N2 K lineage, comprising nine HA substitutions relative to its J.2 ancestor, offers a definitive stress test for predictive models of protein adaptation (Dee et al., 2026). Using this lineage as a case study, we investigate how evolutionary forecasting can be strengthened across different contexts by explicitly controlling for mutation biases. We demonstrate that mutational signatures are embedded in foundation PLMs trained on all proteins (ESM-2 and ESM-C). Critically, we show fine-tuning further strengthens the capturing of the nucleotide mutation bias in the PLM, compounding the impact in downstream applications.

## Results

We first contextualised the H3N2 K lineage defining nonsynonymous mutations by inferring the likely order they happened in and their PLM pseudo-probabilities on the J.2 backbone. The substitution probabilities were generated by embedding the H3N2 J.2 sequence and looking at how likely the model inferred each of the K lineage’s amino acid replacements were on that backbone. All of the J.2 to K lineage defining amino acid replacements involve one nucleotide change per codon and are ranked in the upper regions of probability of the PLM and top third of mutation accessibility (Figure 1B). Additionally, all can be observed in the HA-80 training data to different degrees (all are seen in at least 70 of the 8,099 sequences; though T328A is not observed in human derived training sequences; see Supplementary Table 2). The K lineage amino acid replacement with the highest probability was HA2:S49N with a probability of 0.904 and a rank of 7 (top 0.07%), while the lowest probability change was HA1: T328A with a probability of 4.72×10^-4^ and a rank of 578 (top 5.4%; Figure 1B). All of the J.2 to K inter-lineage amino acid replacements involved a single nucleotide mutation, though they spanned a much larger range of ranks than the PLM: from the 241st most likely mutation (Q173R; top 2.3%) to the 3294th (I160K; top 30.7%; Figure 1C).

**Figure 1.**
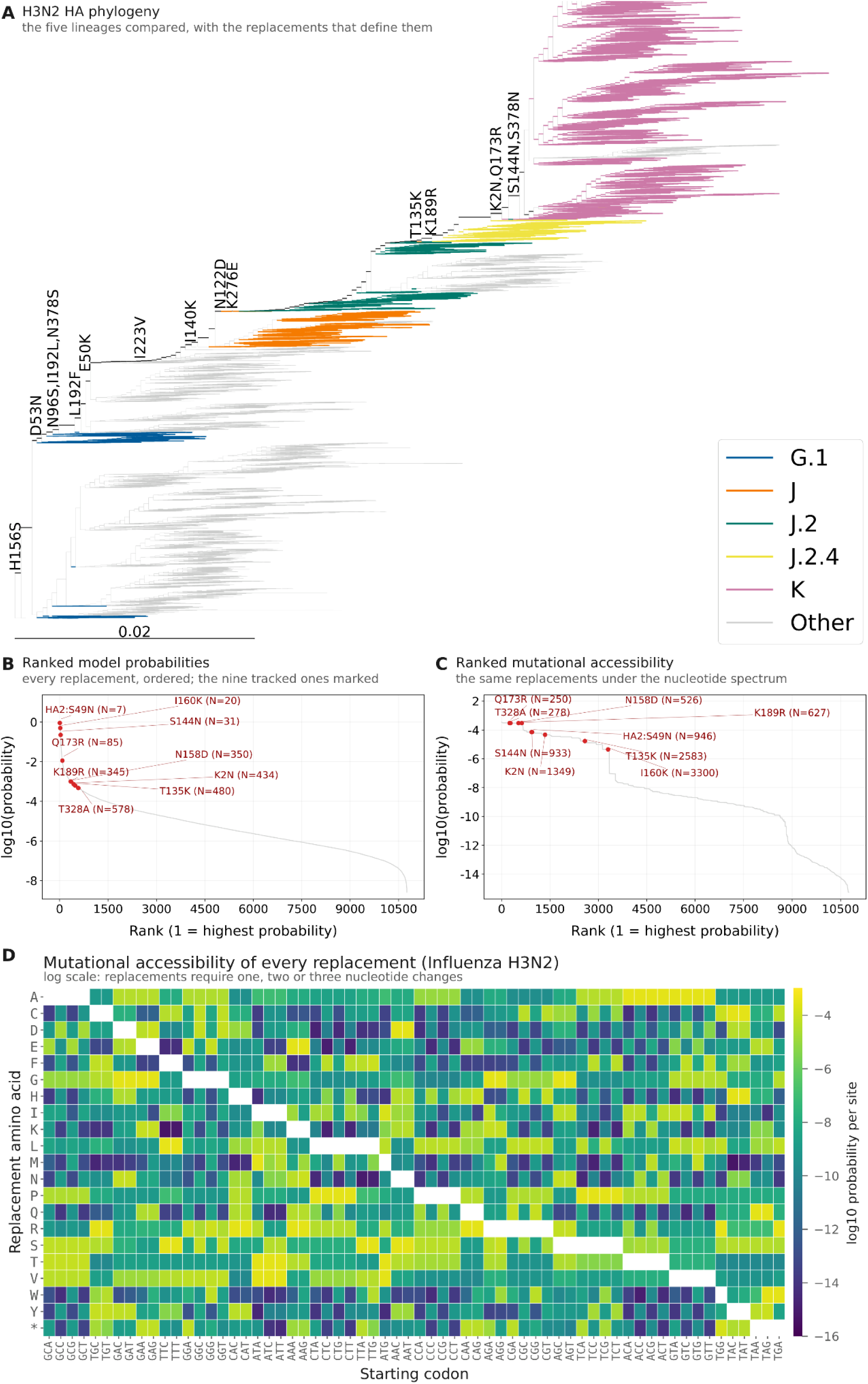
**(A)** Phylogenetic tree depicting influenza A H3N2 K lineage’s emergence (pink), see key for other clade assignment colours. **(B)** Plot of all possible 19×566AA non-reference (HA1 numbering) amino acid replacement probabilities from the HA-80 PLM ranked on the x axis with their PLM probabilities on the y axis (logarithmic scale), the J.2-> K mutated are highlighted along the plot, and their ranks are printed at the end of the point labels. **(C)** Plot of the mutation sets with their log10 mutational probabilities (y) and ranks (x). **(D)** Heatmap of the log10 mutation probabilities to each amino acid from all 64 codons, showing the wide range of possible probabilities.

To test the capacity for PLMs to learn underlying mutational signatures, we investigated how accurately codon-level mutational accessibility can be inferred from the protein sequences alone, i.e., without codon information. We hypothesised that PLMs implicitly learn the mutational forces driving the replacement rates between amino acids. Because different amino acids carry varying amounts of codon-level information—ranging from the perfect information of single-codon variants (Methionine and Tryptophan) to the high uncertainty of six-codon variants (Arginine, Leucine, and Serine)—we sought to quantify the absolute ceiling of mutational accessibility information that can be recovered from amino acid data alone.

To model this informational constraint, we designed a matrix compression and reconstruction pipeline. We first applied this to the empirical H3N2 influenza mutation rates from Pauly et al. (2017) to simulate the species-specific fine-tuning process described by Lytras et al. (2026). The influenza 64×64 codon-to-codon mutation probability matrix was compressed into a 64×20 codon-by-amino-acid matrix, summing over all synonymous target codons for each amino acid (Figure 1D). This was then compressed to a 20×20 amino-acid matrix by averaging across synonymous source codons, effectively assuming equal weighting of synonymous codons. Finally, we reconstructed a 64×20 codon-by-amino-acid matrix by assigning each codon the mean row of its encoded amino acid, and quantified the information lost in compression by comparing the original and reconstructed 64×20 matrices. We report the explained variance score, 100 × (1 − Var(t − p) / Var(t)): the percentage of the target’s variance (t) accounted for by the predictor (p). To represent the unaligned pan-species data on which base models like ESM-2 are trained, we repeated this entire mapping process using a generalised Kimura 80 (K80) one parameter mutation model (Kimura 1980). A panel of other mutation spectra from diverse species were also compared, which all had lower correlations to H3N2’s directional spectrum than K80 (Figure S1).

This comparison revealed 84.8% of the variation in the 64×20 H3N2 specific matrix was retained by compression into the 20×20 matrix. However only 43.6% of this influenza specific variation is captured in the compressed matrix generated under the generalised Kimura 80 mutation model. This stark variance gap demonstrates how much specific information is theoretically available during fine-tuning on a specific organism with no codon knowledge. We assumed uniform codon usage when reconstructing our codon-level mutational profiles, but with fine-tuning on vast quantities of data the PLM could potentially implicitly learn more than this with non-random codon usage. Depending on the clustering methods for fine-tuning, fine-grained intra-lineage diversity may allow a fine-tuned PLM to capture even more of the mutational variance than our estimate. ESM-2 was trained on sequences clustered at 90% identity (which were sampled from 50% clusters to control for ascertainment bias), and HA-80 was then fine-tuned with 99% identity clusters (with a 2015 training data cut-off date).

We thus quantified the extent to which PLMs have implicitly learned mutational accessibility with and without fine-tuning by correlating mutation probabilities with PLM amino acid probabilities. We found that both the original ESM-2 (650M) model and the ESM-2 HA-80 model (650M; fine-tuned on all influenza A HA sampled until February 2015) correlated with mutation probabilities. While mutational factors account for negligible variance in raw PLM probabilities (Pearson r = 0.045 for HA-80 versus 0.054 for base ESM-2), they exhibit much stronger log-scale correlations (r = 0.627 and 0.283 respectively). This shows that while the primary allocation of raw probability mass is driven by factors other than mutational accessibility, the models strongly capture mutational accessibility, particularly after fine-tuning.

We retrained the ESM-2 (650M) PLM as in Lytras et al. (2026) using the same HA dataset to gauge how the mutational accessibility signal was captured throughout the fine-tuning process. The model rapidly learned the accessibility space, plateauing after the second epoch (Figure 2C). We then validated that the strengthening of the mutational signatures in the PLM was not an influenza or ESM-2 unique phenomenon, by also fine-tuning the ESM-C 600M model on SARS-CoV-2 spike protein (using a tweaked training methodology; see supplementary information). The Spearman’s ρ coefficient of ESM-C PLM probabilities to mutational probabilities on this SARS-CoV-2 data rose from 0.299 before fine-tuning to 0.679 afterwards (Figure S2). We found that masking an amino acid did not meaningfully impact the mutational accessibility bias of the PLM pseudo-probabilities (Figure S3).

**Figure 2.**
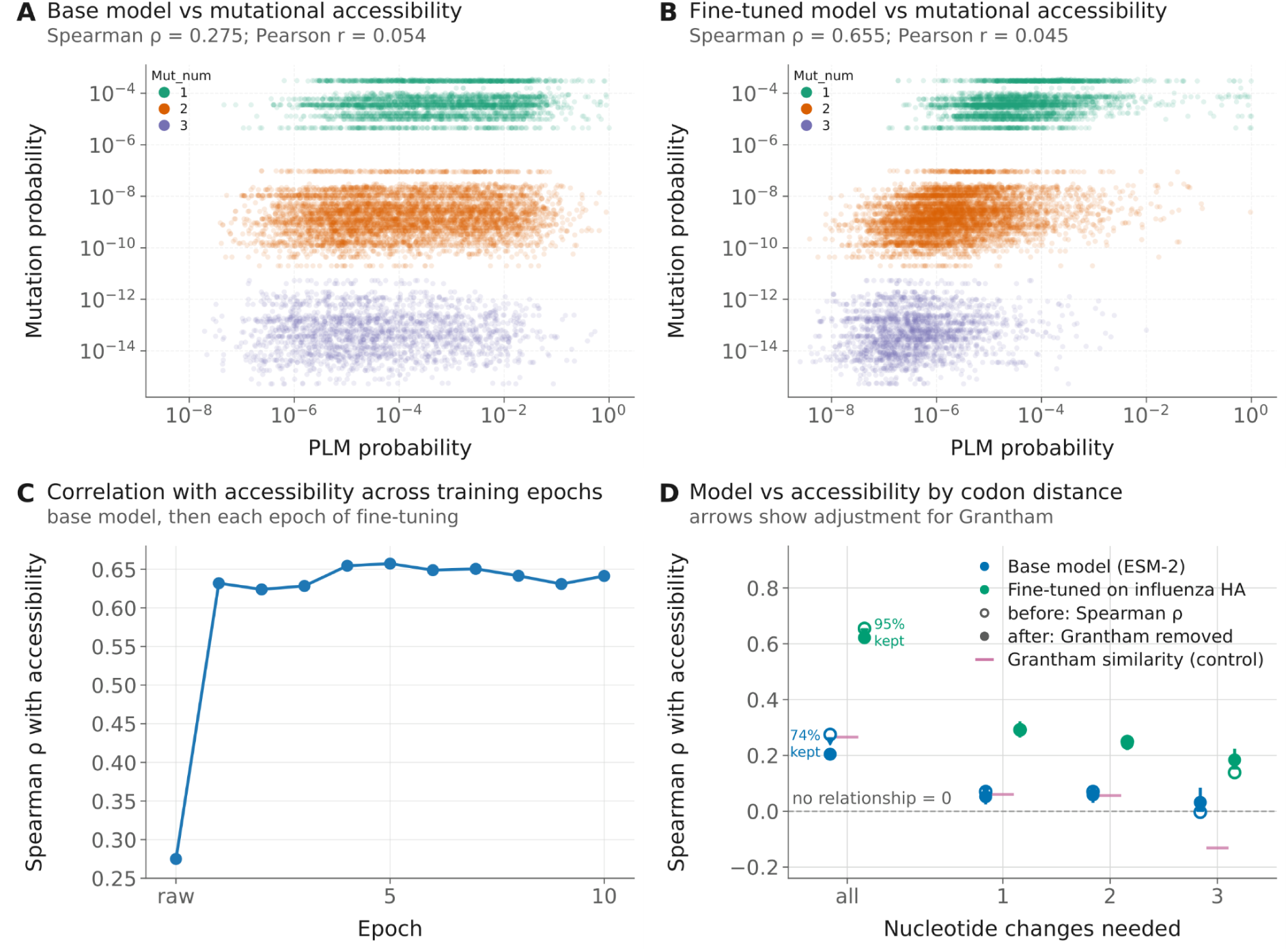
**(A)** PLM probabilities (x axis) for the original ESM-2 (650M) model versus mutation accessibility (y axis) for every possible amino acid change in the 19AA×566 position matrix for the H3N2 J.2 lineage HA sequence. **(B)** PLM probabilities (x axis) for the fine-tuned ESM-2 HA-80 of the same architecture versus mutational probabilities (y axis). The three clusters of points along the mutation axis represent 1/2/3 nucleotide mutations to transition from one amino acid to another. The bottom right points represent mutations assigned high likelihood by the PLM but that are highly inaccessible through mutational changes **(C)** Spearman correlation of mutation probabilities versus PLM probabilities across fine-tuning epochs on the HA-80 H3N2 data, rapidly learns mutation space. **(D)** Decomposing mutational accessibility into nucleotide bins to see if the PLM knows which amino acid changes are more likely within strata as well as across them. Grantham’s matrix as a control for amino acid properties, shown by itself and used as a control with partial Spearman regression to isolate PLM bias.

We then checked to see if the mutational signature was being learned by the PLMs within, as well as across, the nt=1,2 and 3 strata of codon space. We found the correlations were notably weaker here, although non-zero, suggesting the largest component of accessibility learned by the PLM is the distance between codons (at nt=1, the Spearman ρ was 0.292 for HA-80 and 0.071 for base ESM-2; dropping down to 0.139 for HA-80 and-0.003 for ESM-2 within the nt=3 distance stratum). The fine-tuned model shows this bias across all codon distances, showing it has absorbed mutational biases as well as the codon proximities. The codon table puts biochemically similar codons closer together so we used Grantham’s matrix, which summarises these properties as a control. The base ESM-2 model’s pseudo-probabilities do not correlate notably more to mutational accessibility than Grantham’s matrix alone does. We used partial Spearman regression to investigate if the similar correlation coefficients of Grantham’s matrix and mutational accessibility were the same shared signal, or if they were orthogonal. Most of the mutational accessibility correlation in all categories was retained following this adjustment, 5 % of HA-80’s and 26% of ESM-2’s correlation was lost respectively, however all showed a Spearman ρ above 0 for all but the 3nt band, where the ESM-2 Spearman ρ remained around 0 (Figure 2D). The majority of the variance in mutational accessibility between replacements is driven by the number of mutations required (Figure 1D) and this is detected by both PLMs (Figure 2D). The fine-tuned PLM’s pseudo-probabilities also reflect variance across replacements requiring the same number of mutations as well as across them (Figure 2D), demonstrating knowledge of the nucleotide mutation matrix.

To investigate the species-specific nature of the mutation biases absorbed, we compared a panel of different mutational matrices to see if this effect was H3N2 specific or more general. The mutation bias knowledge was found to be more of a general than specific effect as these correlations were shared with other taxon’s mutation matrices (e.g. Escherichia coli and human; Figure S4). However, a symmetric H3N2 matrix correlated better than the directional one, illustrating the non-directional nature of PLM training: there is no temporal ordering of the training data as it is used to refine model weights during training.

We hypothesised that combining the mutational accessibility of amino acid replacements with their PLM probabilities might improve evolutionary predictions versus raw PLM probabilities. This was investigated by trying to predict the intra-lineage diversity of HA in the recent H3N2 sublineages with the PLM pseudo-probabilities. Given that the PLM clearly had some implicit understanding of mutational accessibility (Figure 2B), there was a concern that a simple product of the two probabilities would over-emphasise the mutational accessibility, by representing it twice. Therefore, we performed a parameter sweep, variably weighting mutational accessibility, raising it to the power of α from-1 to 1 to see what would most improve predictions (an α of 0 represents the raw PLM and a value of 1 represents PLM_prob×mutation_prob; see Equation 1).

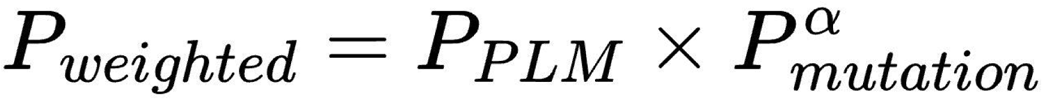

Equation 1: weighting PLM probabilities by the mutational accessibility of the AA replacements (P_mutation_ is calculated by summing the mutational probabilities from a given codon to each of the target amino acid’s synonymous codons)

Predicting future diversity can be broken down into two tasks: 1) which variants will be observed, and 2) which of those variants will have spread most. Looking at the frequency of all possible mutations tests both of those simultaneously. Given the strength of constraint that mutational supply provides, higher α values (stronger weighting of mutational accessibility) would be expected to better predict which amino acid replacements are observed. However, predicting what frequency these replacements rise to may be better predicted by the PLM with the mutational accessibility information removed (a negative α, downweighting more accessible replacements in proportion to a fractional power of their mutational accessibility probabilities) may provide an estimate of protein fitness that is less confounded by mutational accessibility. To test this, the frequency of all amino acid replacements was calculated in the HA protein for 13 lineages between the HA-80 training cut off and up to and including the emergence of the K lineage (filtering out lineage intermediates to avoid cross lineage contamination).

Initial evaluation focused on a combined prediction task: analysing the frequency in the population of all possible amino acid replacements with zeroes included. The fine-tuned model displayed minimal gains in prediction accuracy when the probabilities were multiplied by the mutational accessibility information; Spearman’s ρ shifted from 0.297 (α=0) to 0.313 (α=1; Figure 3A). The mutation accessibility probabilities alone performed similarly with a Spearman’s ρ of 0.285. The base ESM-2 model underperformed both with a Spearman’s ρ of 0.201, though the gap closed to the fine-tuned model when multiplied by mutation accessibility probabilities, rising to 0.299 at α=1.

**Figure 3.**
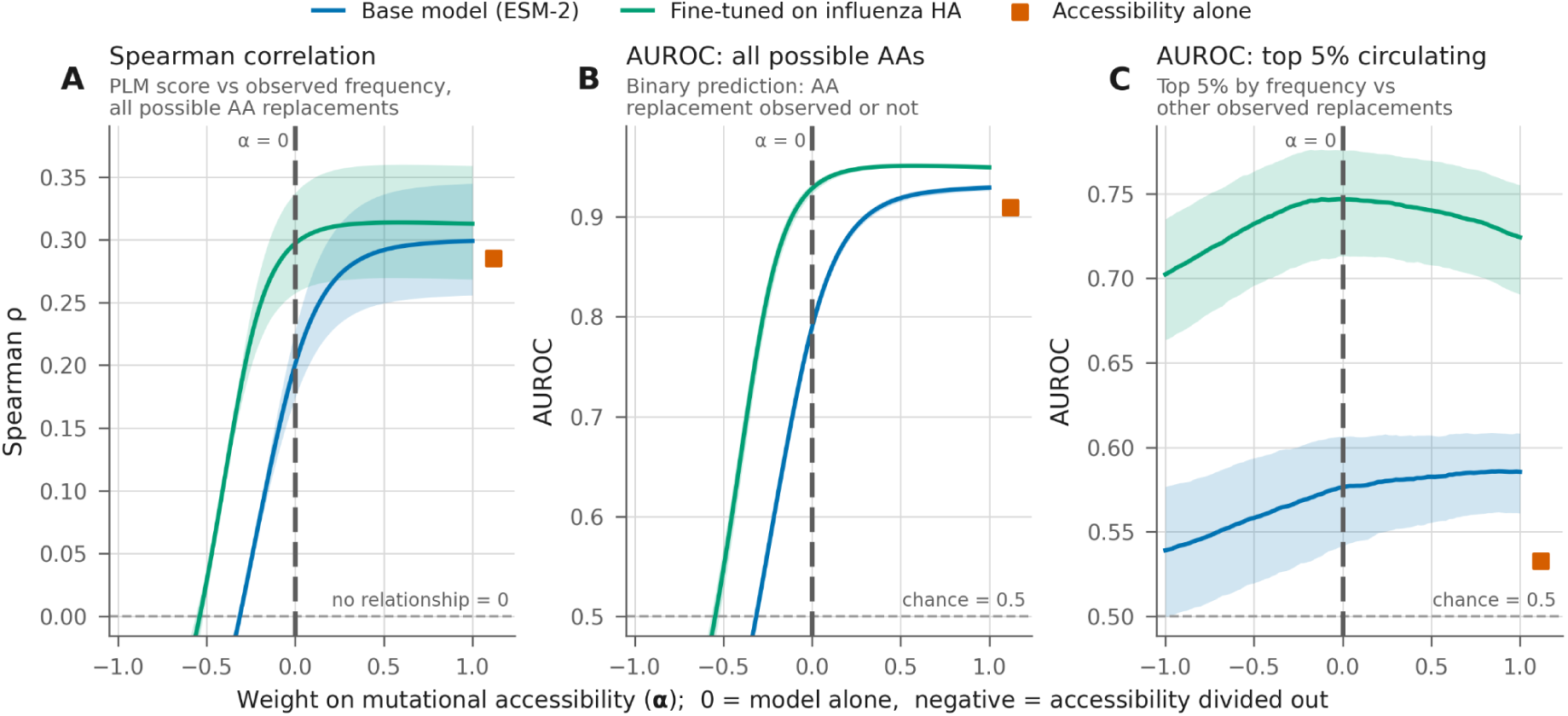
Predicting circulating variation in IAV HA in H3N2 lineages. All values show an average of 13 independent sequence sets from post 2015 lineages (see Supplementary Table 1). The x axis shows a range of α weighting for weighting the PLM pseudo probabilities of mutations by their mutational accessibility. Score=PLM_prob×Mutation_prob^α^. An α value of 0 is highlighted by the dashed line and shows the correlation of the pure PLM probabilities without any explicit mutation accessibility probabilities. **(A)** Spearman regression on all possible non-reference mutations, with non-mutated sites with a frequency of 0 included. **(B)** AUROC for logistic regression for a binary classifier predicting which amino acid replacements are or are not observed in each lineage from PLM scores. **(C)** AUROC for classifying which amino acid replacements are or are not in the top 5% frequency of observed changes in each lineage.

To decompose the evolutionary predictions further, the model was evaluated on its binary prediction performance for which HA amino acid replacements are present and circulating in the 13 major H3N2 lineages circulating after the HA-80 model’s training cut off. The fine-tuned HA-80 PLM (Figure 3B; AUROC 0.928) matched the raw mutation accessibility (0.909) and outperformed the base ESM-2 model (0.789). Incorporating the mutational accessibility probabilities further improved prediction performance for both the fine-tuned and base ESM-2 PLM providing the strongest predictive power (AUROC values rising to 0.950 and 0.929 respectively). These results were robust to a range of enrichment metrics (Supplementary Figure S5; peak PR-AUC enrichment >10 for alpha weighted fine-tuned model).This highlights how selection and mutation combine to shape the circulating diversity present in influenza and that the PLMs do not fully capture mutation accessibility constraints.

Model performance was then evaluated in predicting the fittest amino acid replacements, by predicting the frequency of only the mutated sites circulating in the global population. We again used AUROC but only on the non-zero frequency amino acid replacements circulating in each lineage, to identify the most successful mutations. We classified high frequency replacements as those in the top 5% of allele frequencies within each lineage, leading to different percentage thresholds in each lineage (Supplementary Table 1). The fine-tuned model outperformed ESM-2 (AUROC of 0.747 versus 0.576; Figure 3C), and incorporating mutational accessibility generally made performance worse, especially for the fine-tuned model. Mutational accessibility by itself had a near random AUROC of 0.533, suggesting that very little of the variance in observed allele frequencies was driven by mutational accessibility. This suggests that natural selection dominates the signal once diversity emerges, although we observed many homoplasies in each lineage, most replacements are seen to only evolve once per lineage (Supplementary Table 1). Subtracting the mutational accessibility (α<0) provided no real gains in predictive power from the PLM. This suggests that although the mutational accessibility information is encoded, it isn’t harming the allele frequency predictions. However, this is being tested only on the diversity which exists, which by definition must have been accessible. Additionally, the highest frequency mutations will be influenced by non-selective processes also (e.g., sampling biases and genetic drift), limiting the maximum possible correlation coefficient. Results from synthetic data containing amino acid replacements which are rare in nature may see differing results with negative α values providing stronger performance than the native PLM performance.

To further characterise how the mutational accessibility information encoded in the PLM outputs could confound protein fitness predictions, we investigated predictive capability in deep mutational scanning (DMS) data. We used both the base ESM-2 and the fine-tuned model to predict a panel of different HA experiments. The impact of fine-tuning on prediction accuracy of DMS experiments is mixed. For predicting cell entry, and cell line reproduction (preference ratio) the fine-tuning reduces the Spearman ρ coefficient, but it is notably raised in sera escape, with modest gains in pH stability prediction also. Perhaps as immune escape dominates the recent selective landscape, this shapes the training data more (see Figure S6 for how the structural accuracy of structural epistatic coupling differs across protein regions). For all traits, the fine-tuned models have the strongest prediction accuracy when their probabilities are inversely weighted by mutational accessibility (peak α ranges from-0.24 to-0.18; versus-0.06 to-0.03 for base ESM-2), suggesting that this confounding does hamper protein fitness predictions. A negative α value removed the performance penalty of fine-tuning for cell entry prediction, but not for the preference ratio (Figure 4A and 4D). The poor performance of the fine-tuned model on the preference ratio dataset is most stark at higher codon distances (Figure S7, panel D).

**Figure 4.**
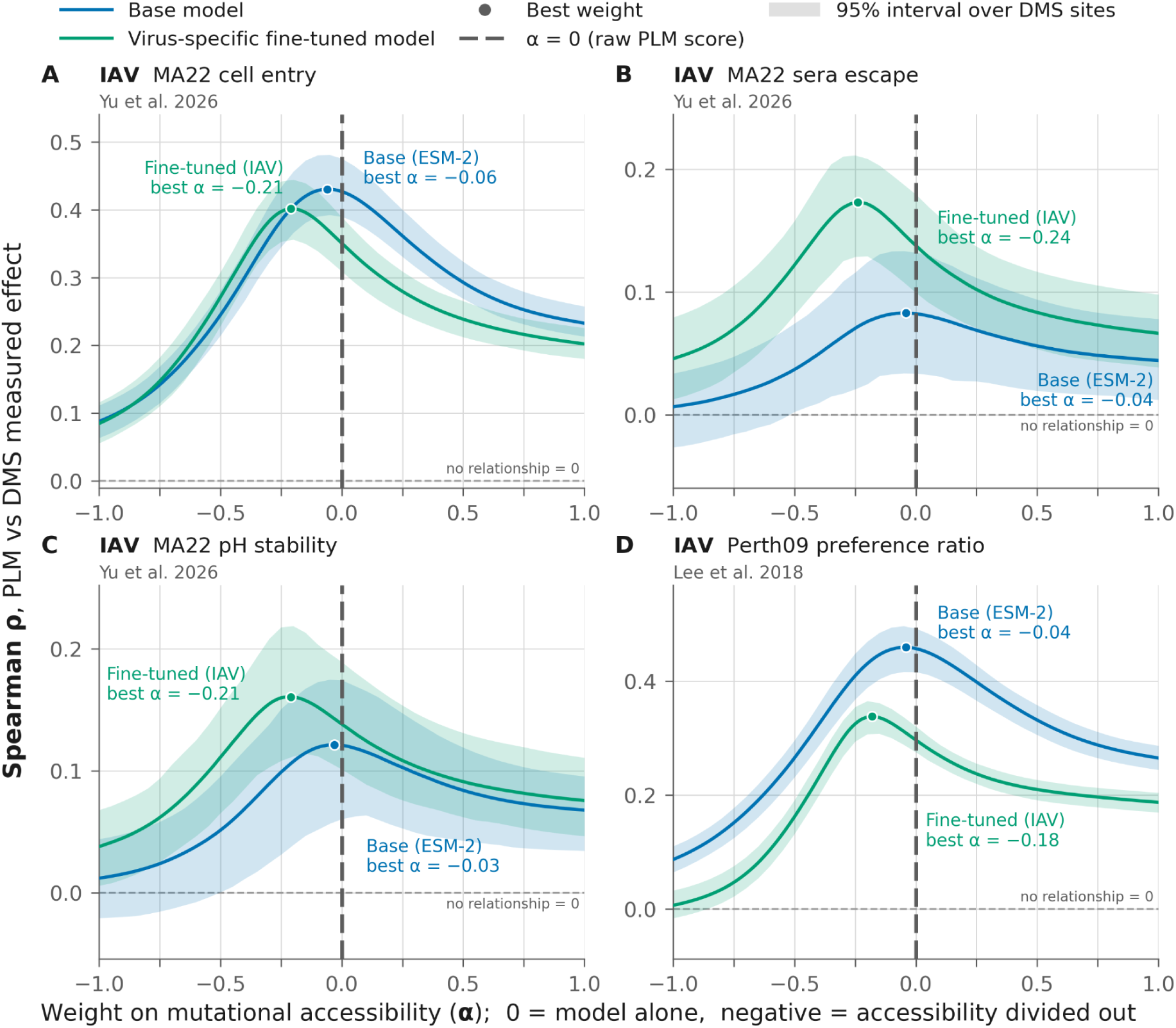
(**A-D)** Deep mutational scanning effect prediction using PLMs for four influenza H3N2 datasets. The x axis shows the weighting α given to mutational accessibility of each possible amino acid alongside the PLM pseudo-probabilities for Spearman rank regression. On all four datasets, a negative α value (discounting replacements by their mutation accessibility) produces the best correlations between the DMS and the fine-tuned PLM scores. The preference ratio in panel D shows a strong advantage of the base ESM-2 model at all α weightings.

We also tested a set of SARS-CoV-2 DMS datasets against that fine-tuned model. The baseline performance of ESM-C was very poor on that dataset, likely reflecting the poor knowledge it seems to have of viral proteins in general (Figure S8), with fine-tuning producing higher absolute Spearman ρ values on all datasets. We also saw the fine-tuned models displayed the strongest correlations with similar negative α values, downweighting PLM scores by the mutational accessibility information (Figure S9). The impact of codon distance on the accuracy of PLM predictions of DMS data differed across datasets. There were significant differences across codon distances for DMS prediction for six of nine datasets(Figure S7). We found negative α values improved fine-tune prediction performance on DMS data for Influenza and SARS-CoV-2 within 1 and 2nt codon strata of codon distances, showing how both elements of mutational accessibility, codon distance and mutation bias can bias PLM predictions (Figure S7).

To further disentangle how protein language models might infer mutational accessibility, we investigated whether they ranked the same amino acid changes as being more or less likely depending on how accessible the starting codon was. For this we only assessed the amino acid pairs which can differ in their accessibility and have at least 5 codons in each of the “easy” (1nt) and “hard” (2-3nt) categories (and at least 15 total codons) in the focal J.2 sequence; this gave 929 “easy” and 1,079 “hard” pair-site evaluations. As the models are trained purely on protein sequences, they should be blind to this; however, the training data is constrained by this accessibility. A codon permutation analysis revealed that the fine-tuned model implicitly absorbs where in codon space each site exists. HA-80 predicted significantly higher average pseudo-probabilities for the same amino acid replacements from more accessible codons (P=0.001; Figure 5A and 5C). The base ESM-C model did not show any detectable directional trends, likely due to the more divergent data it was trained on. A forest plot shows the bias for each individual pair (Figure S10).

**Figure 5.**
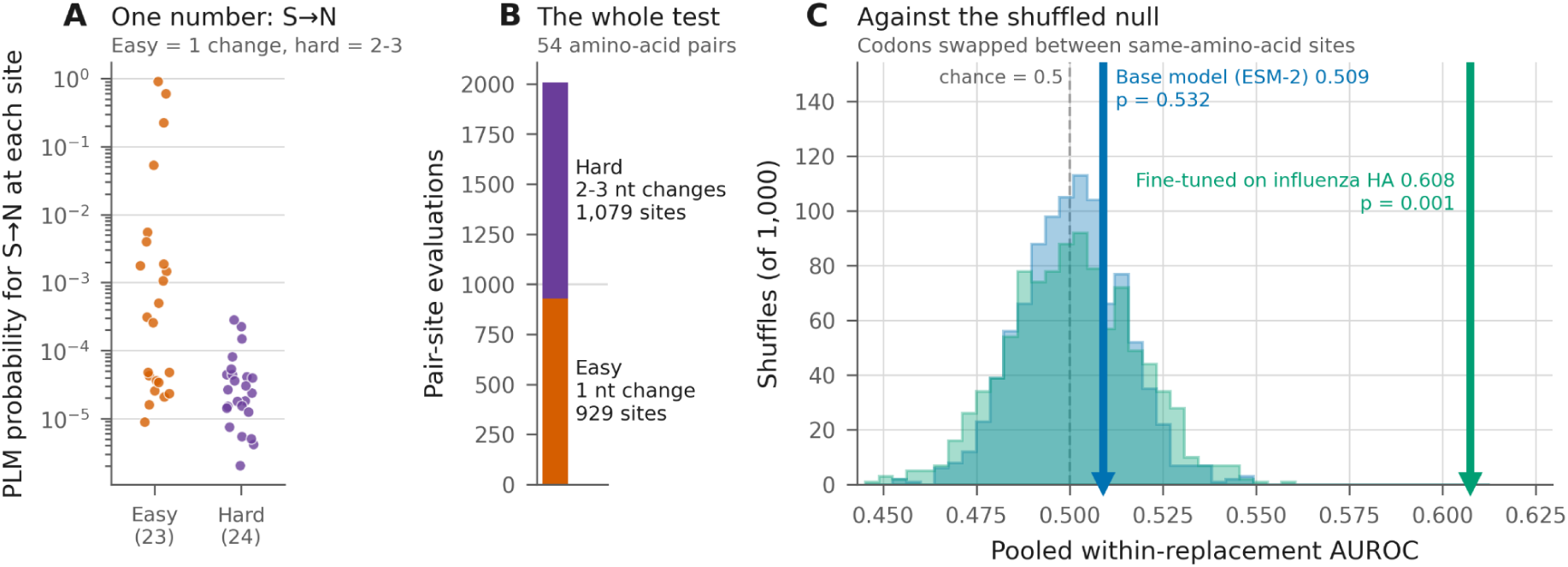
Investigating amino acid substitution pairs which can differ in nucleotide distance in the J.2 ancestral sequence. **(A)** an example directional amino acid pair serine to asparagine can be separated by either one nucleotide or multiple (23 and 24 examples in HA respectively). When encoded in codons requiring only one nt change, the PLM ranks the substitution as more likely than with two. **(B)** showing how many codon pairs fall into each category in HA **(C)** Permutation test across sites, shuffling codons across the sequence to create a null distribution. The fine-tuned model shows a significant non random preference for accessible substitutions in identical biochemical pairs.

## DISCUSSION

PLMs are assumed to map the fitness landscape of proteins; however, our findings indicate they are also learning the underlying mutational accessibility of amino acids in protein sequences. We find that even without seeing the underlying nucleotide sequences, much of the mutational constraint is being inferred by PLMs from amino acid sequences alone. ESM-2 and ESM-C have learned a remarkable amount of mutational accessibility constraints despite training on the amino acid space without explicit nucleotide information. Some mutational accessibility is organism-specific when mutation rates are biased, but some is shared, e.g., when two codons are three mutations apart. Fine-tuning on a specific protein greatly strengthens a PLM’s knowledge of mutational accessibility. Despite influenza’s unusual mutation profile (Figure S1), the fine-tuned PLM’s predictions correlate similarly well with other organism’s divergent spectra (Figure S4). The model appears to have learned the specific codon space of IAV (Figure 5), and a generalised model of mutation trends (e.g. transition to transversion bias). Our simulations support this, with only minor gains on their own spectra versus other spectra, with a flat, even probability mutation spectrum leaving the weakest signal in the fine-tuned PLM and H3N2’s leaving the strongest (Figure S11). This implicit accessibility knowledge results in a composite output that conflates structural protein fitness with mutational probability.

It is important to acknowledge, however, that the underlying nucleotide sequence and mutations therein are also a direct target of natural selection, and can influence fitness. This is mediated by mechanisms such as host-protein hypermutation of dinucleotides which can then bias codon pairs (Greenbaum et al., 2014; Kunec and Osterrieder 2016). Additionally, different synonymous codons can influence translational dynamics, e.g., the utilisation of fast and slow codons, dictating optimal protein folding during translation (Yu et al., 2015; Pinto et al 2018). Therefore, operating exclusively in amino acid space ignores crucial biological constraints. Architectures such as cdsBERT (Hallee et al., 2023) attempt to bridge this gap by shifting the predictive focus from amino acids to specific codons.

These results raise the question of whether fine-tuning a PLM on your specific organism of study improves or degrades prediction performance. For all evolutionary predictive tasks we investigated, fine-tuning improved the predictions (Figure 3). However, multiplying the baseline more naive PLM probabilities with mutation probabilities can produce predictions as accurate as a specialised fine-tuned model, e.g., predicting which amino acid changes will be observed in the population (Figure 3A; 3B). The DMS results show a mixed picture, with certain metrics suffering and others improving, likewise structural inference from epistatic coupling (Figure S6). The initial knowledge of the PLM of the organism in question is likely a key factor, which is very jagged across different viruses (Figure S8), DMS predictions for SARS-CoV-2 improved more consistently than for IAV, perhaps reflecting the apparent ignorance of ESM-C on SARS-CoV-2’s Spike protein. These results (including our simulated datasets using a diverse range of different taxa’s mutation spectra and downstream fine-tuned models; Figure S11) suggest that many organism specific trained/fine-tuned PLMs will have an inbuilt bias towards suggesting higher fitness to more accessible variants. There have been some notable recent efforts in this area (CovFit and DeepCov), which have fine-tuned PLMs with masked learning of a specific virus, alongside incorporating DMS data to optimise the fitness landscape inference (Ito et al., 2025a; Yang et al., 2026). Whilst it’s clear that the fine-tuning can have some detrimental impacts on DMS correlations from PLMs, it remains unclear if grounding the model’s output with DMS data will fully compensate for this bias.

To maximise predictive accuracy of protein fitness using PLMs, there is clear value in explicitly decoupling mutational accessibility from protein fitness and utilising both information types. Ideally, a PLM would evaluate protein structure-mediated fitness entirely independently of mutational constraints. This isolated metric would be highly valuable for synthetic biology, where mutational accessibility is rarely a limiting factor. However, synthetic libraries will have their own biases; for instance, deep mutational scanning libraries often utilise restricted degenerate codon spaces (e.g., NNK codon libraries). Fine-tuned PLMs may implicitly learn these experimental distributions leading to suboptimal protein functional predictions. For evolutionary forecasting, researchers could then model trajectories by explicitly combining predicted fitness with mutational accessibility. This would provide more meaningful predictions than the aggregated metrics that current state of the art models provide. Matsen et al. (2026) have developed such an approach using phylogenetic sequence pairs, by predicting child sequences from parental ones with a defined mutation probability matrix. They successfully applied this method to antibody sequences, where the limited diversity of non-mutated antibody sequences results in particularly strong mutational accessibility biases.

We mostly think of fitness valleys as being shaped by epistatic constraints, requiring multiple amino acid changes to cross a region of low fitness. However, assessing 30,028 individual amino acid replacements across the 50,094 internal branches with mutations in the H3N2 Haemagglutinin phylogeny, only six amino acid replacements requiring three nucleotide changes are observed on single branches. This compares to 29,875 amino acid replacements requiring one nucleotide change along single branches, and 147 requiring two nucleotide mutations (Figure S12). This suggests the evolutionary accessibility of certain point amino acid replacements is likely to require viable intermediate nonsynonymous codons to traverse codon space in an analogous fashion to inter-site epistasis. These fitness valleys limit the training data for PLMs to be able to predict distant codon space accurately, this is observed as DMS prediction accuracy dropped off for certain datasets at nt=3 codon distances (Figure S7).

Ultimately, the vast combinatorial and stochastic nature of possible mutations mean that viral evolution will never be entirely predictable. Nevertheless, explicitly separating pure protein fitness from mutational accessibility biasing amino acid changes and direct selection at the nucleotide level (dinucleotides, fast/slow codons for translation etc.) will likely yield the most robust predictions. This is important for both evolutionary forecasting where mutational supply is a crucial factor, and to optimise synthetic systems free from evolutionary history and constraints. As we have demonstrated here, accounting for the distinct contributions of mutational supply and structural selection will offer a clearer framework for accurately mapping and forecasting real-world evolutionary trajectories.

## Materials and Methods

### Sequence data for diversity analysis

Sequences for influenza A were obtained from Genbank using our viral genome toolkit software, code available at: https://github.com/centre-for-virus-research/V-gTK (DOI:10.5281/zenodo.23504636), and supplemented with GISAID records (up to 12 February 2026). Sequences were restricted to H3 subtype HA coding sequences. (GISAID acknowledgements in Supplementary Section 2).

### Analysis code

All analysis code outwith the v-gTK functionality, and required intermediate tables has been deposited in its own github repository https://github.com/omaclean/PLM_mutational_accessibility (DOI: 10.5281/zenodo.23889217). All figures should be re-generatable from there.

### Diversity analysis

For the intra-lineage diversity analysis we used the precursor lineages C.1, D, E, E.2, G.1, G.1.1, G.1.3, G.2, G.2.2, then split the J to K lineage transitions up as: J, J.2, J.2.4, and K for a total of 13 lineages. We took the Nextclade/Nextstrain H3N2 subclade definitions as the root sequence for each lineage from https://github.com/nextstrain/nextclade_data/tree/master/data/nextstrain/flu/h3n2/ha/EPI185 7216. We defined a sequence as belonging to that lineage if it had all of the lineage defining amino acid replacements and no more than five additional amino acid replacements from the Nextstrain ancestral maximum likelihood lineage root. We used a stringent filtering approach to keep our lineage data separate and avoid intermediates between the named lineages. We filtered out any sequence which possessed more than two of the lineage defining amino acids of the next descendent lineage(s). After filtering, this left: 86,661 sequences in total, each lineage’s sample size ranged from 776 J.2.4 sequences to 26,703 J.2 sequences. A detailed metadata summary table can be found in Supplementary Table 1. Amino acid replacement frequencies were calculated as a proportion of the number of available sequences. Each lineage analysis contains independent sequence sets, therefore because HA does not recombine, each lineage’s diversity is an independent data point. Intervals are bootstrapped across lineages treating them as 13 replicates not 13 measurements of a single population.

### Phylogenetics

The influenza phylogeny for discovering mutation ordering followed the methodology of Dee et al. (2026). Briefly: influenza A virus sequences were retrieved from the NCBI and supplemented with GISAID data. Sequences were restricted to H3 subtype HA coding sequences from human hosts. Records failing quality control were excluded: HA length < 1680 nucleotides (excluding N and non-IUPAC characters), atypical genetic divergence, or inconsistent host annotations. Redundant records were identified by strain name and removed to minimize duplication across databases. Sequences were clustered with MMseqs2 (v15.6f452; Steinegger & Söding 2017) at a 0.988 sequence identity threshold. Cluster representatives sequences were aligned using MAFFT (v7.453; Katoh & Standley 2013) with default parameters and a maximum-likelihood phylogeny was inferred with IQ-TREE (v2.4.0; Minh et al., 2020) under the best-fit substitution model (GTR+G+I). IQ-TREE was used to reconstruct a comprehensive tree for the G.1 clade and descendant lineages. Finally treetime (v0.10.0; Sagulenko et al., 2018) was used to reconstruct the ancestral amino acid changes along the backbone of the phylogeny.

### Mutation accessibility

Following the methodology of Gunnarsson and Babu (2023), mutation accessibility calculations between codons were calculated for a single round of replication. They use the observed mutation probabilities from Pauly et al. (2017) per replication round of H3N2. They defined variant accessibility as their probability of being gained through one round of replication. Host led mutational processes on the positive stranded intermediates drive a strong directional bias in H3N2 creating an unusual spectrum. For example, the codon mutation probability of AAA->GGA (9×10^-8^; Lys->Gly) is the square of the probability of AAA->GAA (3×10^-4^; Lys->Glu) in our model, as the probability of the double mutant is the square of the single mutant. For target amino acids with multiple synonymous codons, we sum over the probabilities of reaching any of the target amino acid codons. We consider only direct substitutions between codons, excluding pathways involving intermediate codons; stop codons are excluded from PLM comparisons.

### Mutation matrix compression

We collapsed our 64×64 codon matrices to 64×20 codon x amino acid matrices by taking all synonymous codons as equally valid targets and summing their probabilities. To compress the 64×20 to a 20×20 transition matrix, we assumed equal codon usage. We suspected that the small nature of the HA protein would lead to very noisy estimates of biased codon usage. We reconstructed a 64×20 codon-by-amino-acid matrix by assigning each codon the mean row of its encoded amino acid. We quantified the information loss in the compression by converting the matrix to a list of probabilities and comparing the variance of the original matrix retained after compression using Nash–Sutcliffe model efficiency. We also tested how a more generally trained model using a more neutral set of mutation assumptions would perform. To do this we calculated 64×64 codon matrices using a simplified model which allows for equal base probabilities but with a biased transition to transversion ratio, for which we used a value of 2 (Kimura 1980), and repeated the above process.

### Protein embedding

To explore the grammaticality of sequence changes we used the method from Lamb et al. (2026). We embed an unmasked sequence in a single forward pass in the ESM-2 framework and extract the model’s amino acid pseudo-probabilities in a 20×566 amino acid matrix. The probabilities are formed from a softmax function on the last embedding layer. We use a single forward unmasked pass for assessing baseline mutational accessibility bias. However for allele frequency forecasting and DMS inferences, we use iterative embedding, masking each amino acid sequentially, and calculate the probabilities the same way from the 20 amino acid tokens at each site.

Our fine-tuned Haemagglutinin models came from Lytras et al. (2026), we used the HA-80 model on the ESM-2 framework to A) get a direct comparison on the same architecture of ESM-2 with the baseline model before fine-tuning and B) because its 2015 training cut off precedes J or K lineages to avoid potential over fitting on the observed J defining mutations.

### Epistasis calculation

We investigated epistasis following the categorical Jacobian methodology of Zhang et al. (2024) and Lamb et al. (2026) and looking at how the PLM’s mutation probabilities shifted as we embedded the unmutated and mutated full length HA sequences in the PLM (Figure S13). We found the zerosum method of Zhang et al. (2024; see equation below) gave the best structural predictions. It operates in log space, and subtracts each position’s marginal average across all 20 amino acids. Note that this can cause some sign changes, where a secondary replacement is increased in raw probability (indicating positive epistasis), and the epistatic coefficient becomes negative after the transformation (indicating negative epistasis).

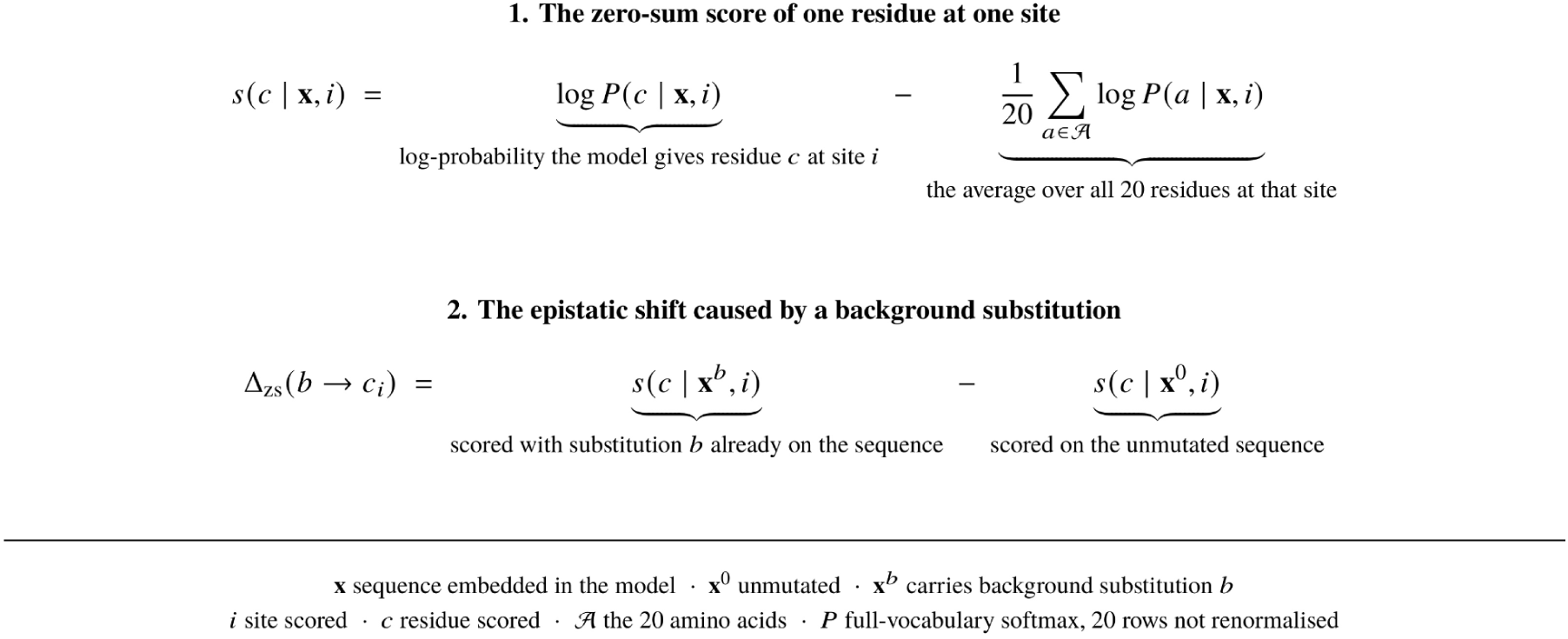

### Grantham’s matrix controls

Because codon space places similar amino acids closer together, we sought to determine whether the PLMs were simply exploiting mutational accessibility as a confounding factor from biochemical similarity. For each possible lineage J.2 replacement we took three quantities: the model’s probability (m), mutational accessibility (a) and the Grantham control (c). We used Grantham as it does not derive from evolutionary data and therefore is not confounded by mutational accessibility, in contrast to BLOSUM62 which is a matrix generated by both processes. We replaced each value by its rank, and computed the partial correlation between ranked m and ranked a controlling for ranked c; i.e. the Pearson correlation between the residuals of ranked m and ranked a after each is regressed on ranked c (Figure 2D; Figure S14, panels A and B). We do these ranks for all possible amino acid replacements pooled across sites, and again within each nucleotide distance stratum. We calculate CIs by bootstrapping across sites.

Because a partial correlation removes only the linear in ranks part of the control’s effect, we also tested two additional supplementary analyses: (i) a Grantham-matched Spearman, in which replacements are binned into quintiles of Grantham distance, the correlation is computed within bins and pooled on Fisher’s z scale weighted by n-3 for the coefficient (Figure S14 panels A and B). We additionally (ii) tested this on a within-site softmax fit to double check that we weren’t observing a methodological artifact of ranks (Figure S14 panels C and D). We found that the base model showed minimal to no correlation within individual strata, only at the global pooled level. The nt3 stratum showed the weakest correlation in all methods, falling to being negative in the base model (Figure S14 panels C and D). CIs were calculated for both metrics by bootstrapping across sites.

### PLM mutation accessibility combined sweep

We swept different weightings of mutational accessibility by sweeping a parameter α to which the mutation accessibility probabilities were raised. An α of 0 means the probabilities were the raw PLM probabilities. Our parameter sweep for combining mutational and PLM probabilities was performed on each lineage separately and aggregated by taking the mean of the correlation values across lineages. We used three metrics, a Spearman rank correlation on the combined task of predicting which sites would be mutated and what frequency they would be. An Area Under the Receiver Operator Curve (AUROC) which measures the accuracy of predictions allowing for imbalanced classes to predict which sites would show diversity in each lineage. Alternative metrics were tested for consistency in Figure S5. Then finally an AUROC prediction only the amino acid replacements with non-zero counts in each lineage, to assess how accurately the models can predict which amino acid replacements are likely to spread best. AUROC was measured on logistic regression classifying the top 5% highest frequency replacements versus the bottom 95%. Code is in the main repository for the paper at: https://github.com/omaclean/PLM_mutational_accessibility/blob/main/evotune_gradient_MAGMA_ESMC.py

### DMS investigation

We correlated the pseudo-probabilities from the PLM versus the observed values from four H3N2 Haemagglutinin DMS dataset (Lee et al., 2018 and Yu et al., 2026). We followed the same α sweep methodology as for the diversity prediction (see Figure S9 for SARS-CoV-2 datasets). The datasets pertain to cell entry, pH stability, sera escape, and preference ratio which reflects in cell growth rates. We embed the sequence of the lab strain used in each assay for each plot as we find DMS dataset concordance decays with genetic distance (Figure S15). We used masked token embedding for each amino acid, finding that in line with Allman et al. (2025) this improved the fine-tuned model’s performance, also seeing it did not notably impact the mutational accessibility bias (Figure S3).

### SARS-CoV-2 fine-tuning

We used the same fine-tuning method as described in Lytras et al. (2026). Briefly, we clustered sequences down to the 99% level with MMseqs2 (Steinegger & Söding 2017) and randomly masked 15% of amino acid tokens in each sequence, calculating entropy loss over these masked tokens. However, we used the ESM-C 600M model instead of ESM-2. We also implemented the MuonW optimiser and MAGMA (Liu et al., 2025; Joo et al., 2026) to improve training speed and optimisation as they were found to cause faster convergence with reduced loss (code is available on github at https://github.com/omaclean/PLM_mutational_accessibility/blob/main/evotune_gradient_MA GMA_ESMC.py).

To adjust for SARS-CoV-2’s distinct mutation profile – which is driven by host proteins acting on the opposite sense RNA genome to influenza A’s genome (Graudenzi et al., 2021) – we used the relative 4×4 nucleotide mutation rates from Table 1 in De Maio et al. (2021). We then normalised these by a 1.3 × 10^-6^ mutations per base per replication rate from Amicone et al., 2022 and adjusted for the SARS-CoV-2 Hu-1 reference genome composition (29.9% A, 18.4% C, 19.6% G, 32.1% U) giving a scaling factor of 8.19 × 10^-7^.

### Codon usage permutation analysis

We used the J.2 root sequence and found all amino acid pairs which differ in the number of nucleotides between them across different synonymous codons. We stratified the synonymous codons for each amino acid pair by their mutational distance, classifying starting codons requiring a single-nucleotide mutation as “easy” and those requiring multi-nucleotide mutations as “hard”. We then compared how the PLM rated identical amino acid replacements from those different starting positions to test if the PLM was distinguishing between the underlying synonymous codons for its predictions. We permuted these codons across sites to form a null distribution of AUROC for each PLM, finding slightly different nulls for base ESM-2 and HA-80 (Figure 5).

## Data Availability

All raw sequence accessions and associated metadata are accessible via the GISAID EPI_SET identifier EPI_SET_260501cz (DOI: 10.55876/gis8.260501cz). Python analysis scripts, trained model checkpoints, intermediate data frames, and plotting pipelines to reproduce all primary and supplementary figures are openly archived in Zenodo at DOI:10.5281/zenodo.23889217 and available on GitHub at https://github.com/omaclean/PLM_mutational_accessibility.

## Supporting information

full supplementary section

## Acknowledgements

We gratefully acknowledge all data contributors, i.e., the Authors and their Originating laboratories responsible for obtaining the specimens, and their Submitting laboratories for generating the genetic sequence and metadata and sharing via the GISAID Initiative, on which this research is based. The CVR authors acknowledge funding from the UK Medical Research Council (MRC, MC_UU_00034/5, MC_UU_00034/6) and UKRI project FluTrailMap-One Health (MR/Y03368X/1). KY and DLR acknowledge funding from the Biotechnology and Biological Sciences Research Council (BBSRC, BB/V016067/1), and KY acknowledges CRUK Scotland Institute (A31287).

