## Supplementary material for "Protein language models learn the underlying accessibility of mutations alongside protein fitness landscapes": full supplementary section

### Supplementary Figures

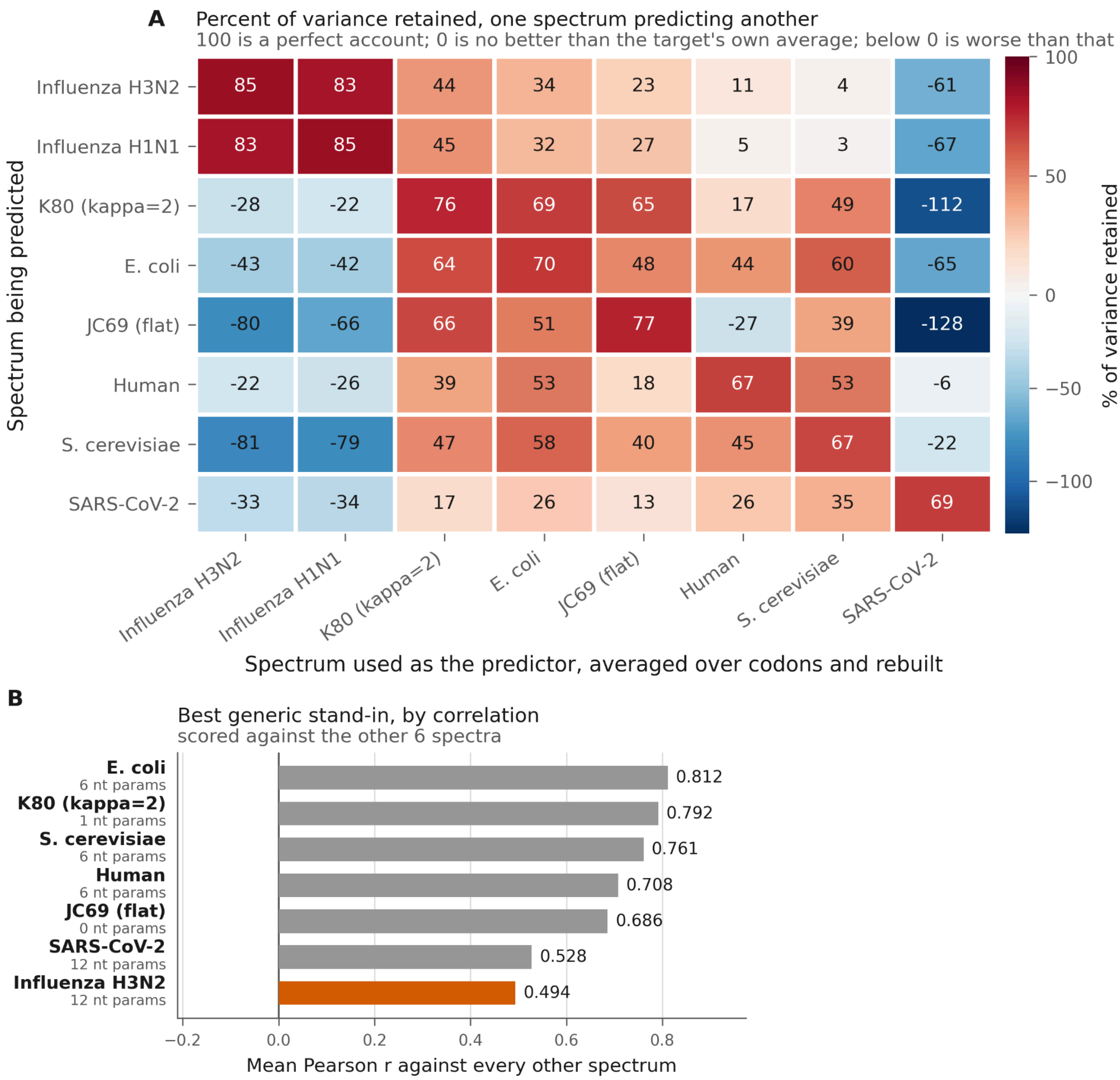


**Figure S1** **(A)** % of variance retained, estimating using Nash–Sutcliffe model efficiency, during a 64x20 amino acid accessibility matrix compression to a 20×20 matrix for different mutation matrices. **(B)** rating generic performance of each spectrum on the other spectra (ignoring H1N1 to avoid double counting influenza).

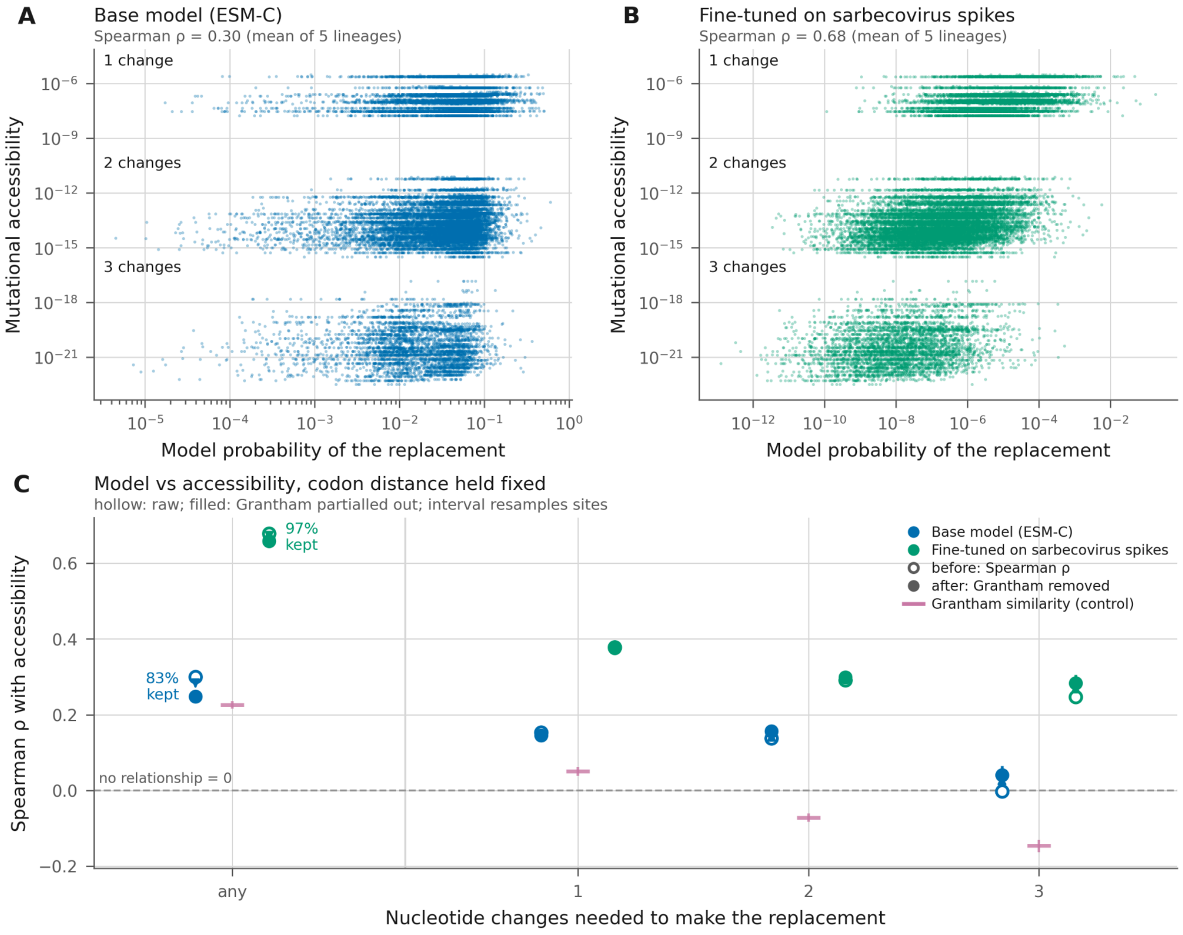


<u>**Figure S2**</u> **(A)** PLM probabilities (x axis) for the original ESM-C (600M) model versus mutation accessibility (y axis) for every possible amino acid change in the 19AA×1273 position matrix for the SARS-CoV-2 Spike sequence. **(B)** PLM probabilities (x axis) for the fine-tuned model on SARS-CoV-2 and wider coronavirus data of the same architecture versus mutational probabilities (y axis). Spearman ρ- 0.299 before fine-tuning to 0.679 afterwards **(C)** Spearman ρ correlation of PLM probabilities with mutation probabilities at different minimum codon distances. Grantham's matrix correlation shown and controlled for.

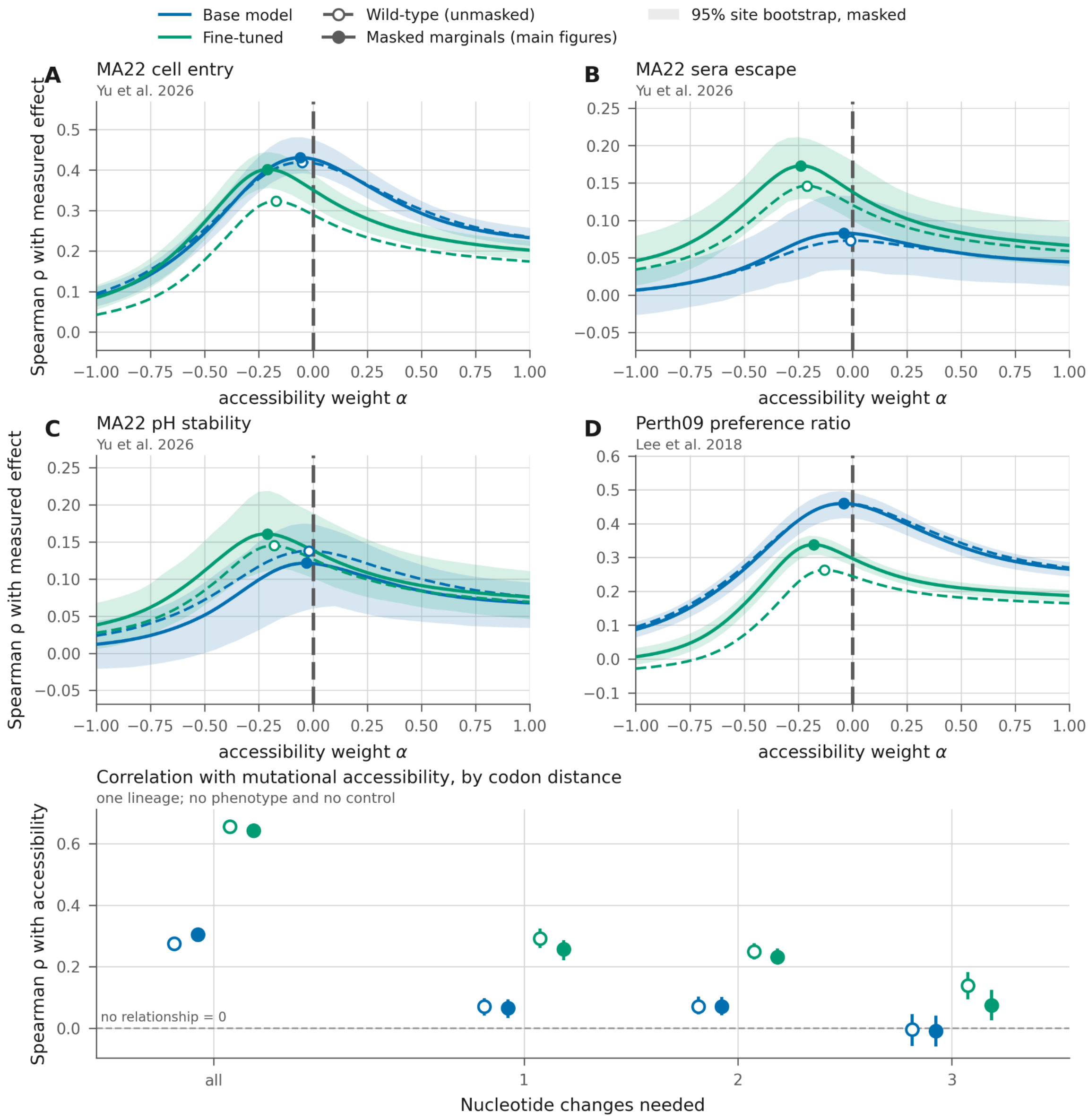


**Figure S3** Allman et al. (2025) show that masking can improve performance for a given amino acid over a single forward unmasked pass. We test this by masking each site for each of the DMS base sequences and see how the new pseudo-probabilities compare to the DMS dataset. **(A-D)** Comparison of DMS prediction accuracy for Figure 4 with a single unmasked PLM embedding sweep dashed line versus an interactive site by site masking approach (solid line). We find that it improves the performance of the fine-tuned model, and does not meaningfully shift the impact of mutational accessibility weighting. **(E)** Masked (full dots) versus unmasked (hollow dots) show the Spearman ρ of PLM embeddings versus mutational accessibility. Masking has a very minor impact on the mutational accessibility bias of the PLM predictions, slightly decreasing the bias in the fine-tuned data and slightly increasing it in the base ESM-2 model.

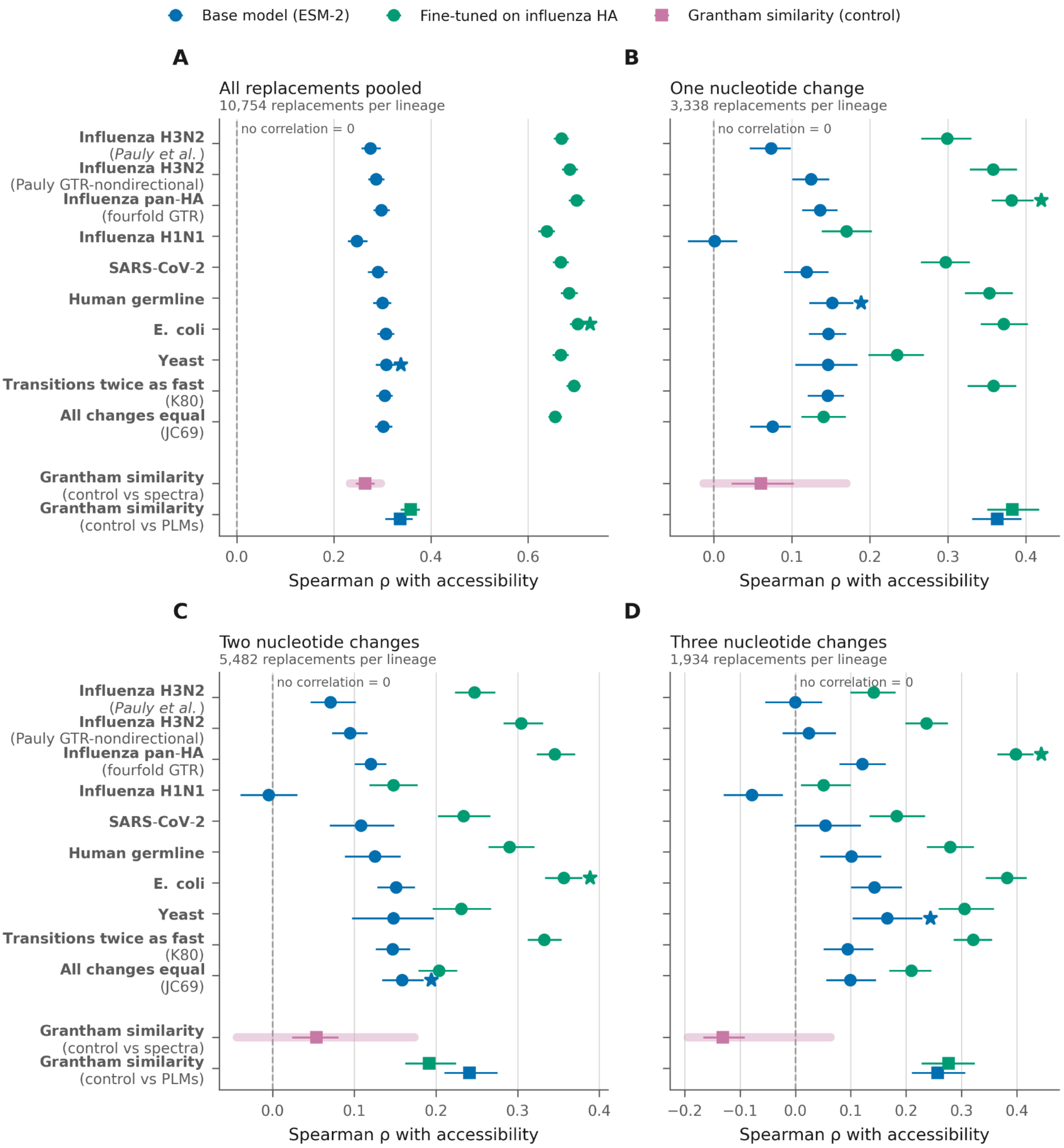


**Figure S4** Correlation of different mutation spectra with base ESM-2 and HA-80 influenza fine-tuned PLMs. The Grantham similarity score is correlated against the H3N2 spectrum from Pauly et al. (2017) for the pink squares, the background band spans the highest and lowest Spearman ρ values against all other plotted spectra. Star symbols show the highest correlating spectra against the PLMs for each codon distance panel. A pan-HA GTR matrix was estimated across the UShER tree from fourfold synonymous sites; it provides a better correlation than the two versions of the Pauly et al. H3N2 directional matrix across the panels. We also plotted Grantham similarity score vs the two PLM as the bottom row for additional context for the relative strength of the mutation bias vs representation of fundamental biochemical properties. **(A)** all possible HA amino acid replacements together **(B)** amino acid changes requiring only one nucleotide mutation **(C)** amino acid changes requiring two nucleotide mutations **(D)** amino acid changes requiring three nucleotide mutations.

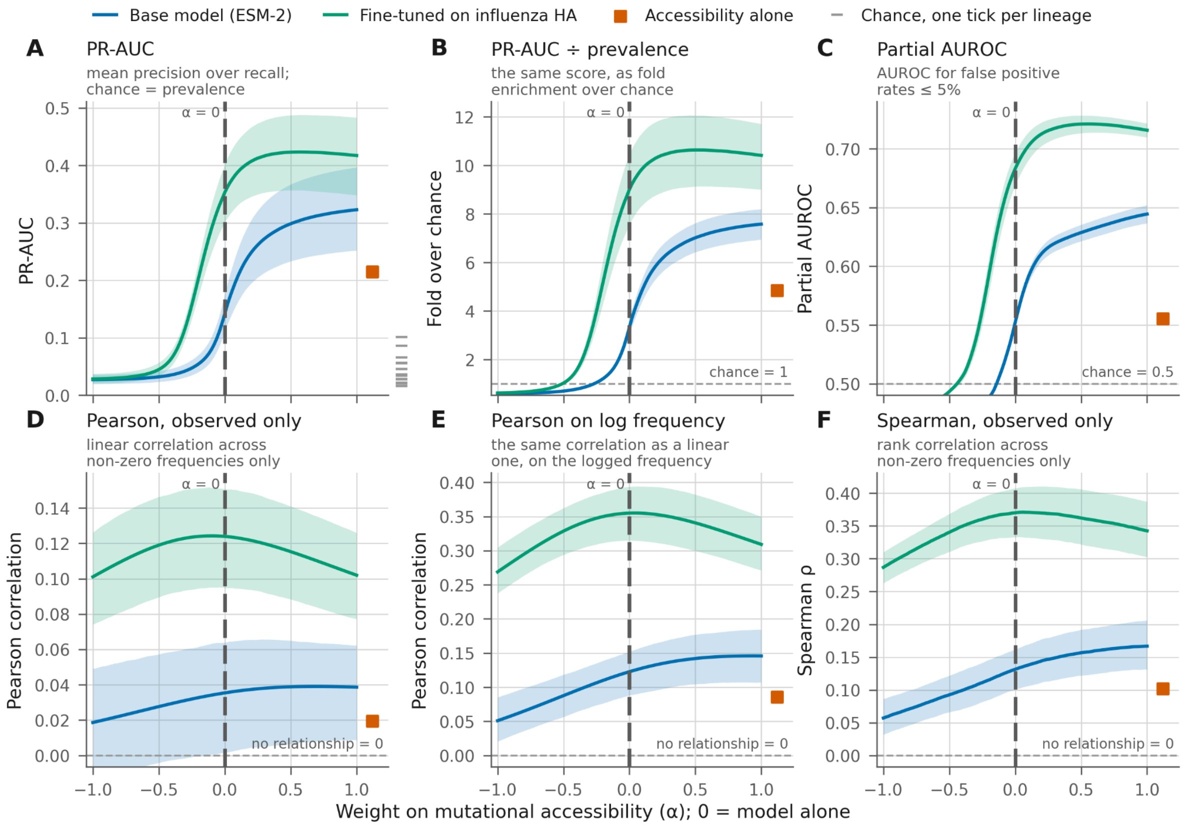


**Figure S5** **(A-C)** alternative metrics for binary prediction of amino acid replacement presence in 13 lineages contrasting to the AUROC metric used in Figure 3B. The observed trends in Figure 3 are robust to all metrics. The dashed lines in the bottom corner of panel A represent the baseline prevalence in each lineage (ranging from 1.6% in lineage J.2.4 to 10.1% in J.2). Panel B adjusts for those baselines. **(D-F)** Alternative metrics for analysing the frequency of circulating variation in the 13 lineages (non-zero frequency amino acid replacements only). **(D)** shows pearson regression on all non-zero allele frequencies, weighting the highest frequency alleles most strongly, increased α weighting on mutational accessibility shows little benefit to the base ESM-2 model, and no benefit to the fine-tuned model **(E)** shows Pearson correlation on logged non-zero frequencies which improves with higher α values, as the mutational accessibility alone out performs the base ESM-2 model, perhaps because of low frequency homoplasies, or because amino acid interchangeability correlates with accessibility. **(F)** shows the Spearman correlation, with a similar trend to the logged pearson score.

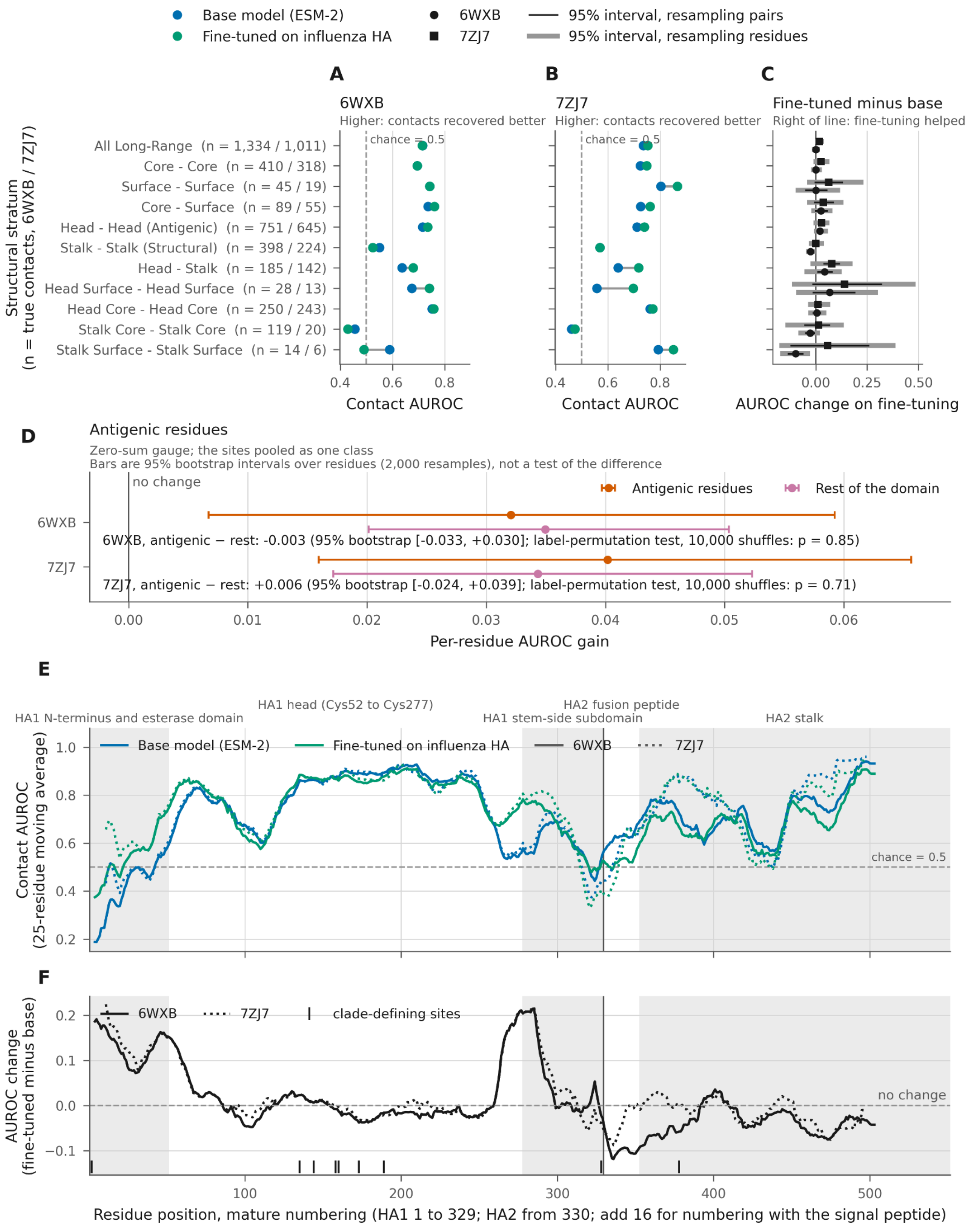


**Figure S6** regions of epistatic coupling performance using Zhang et al. (2024)'s zerosum method using two different HA structures as ground truth. The two structures relate to a cleaved neutral-pH $HA_1/HA_2$ (PDB: 6WXB, pH 7.4; Tan and Rubinstein 2020) and an uncleaved precursor $HA_0$ at acidic fusion pH (PDB: 7ZJ7, pH 4.8; Garcia-Moro et al.,

2022(Tan and Rubinstein 2020; Garcia-Moro et al., 2022). PLM site coupling metrics are used to predict binary structural contact using a threshold contact distance of 8 angstroms and AUROC is calculated. **(A-C)** The structure is split into different regions and fine-tuning (green) is compared to base ESM-2. Fine-tuning improves some regions but generally seems to change very little on average. **(D)** antigenic units show the same marginal gain as non-antigenic regions from fine-tuning. **(E)** The two isoforms (cleaved and uncleaved, and differing pH) of HA have slightly differing performance across regions. **(F)** shows the AUROC difference between fine-tuned HA-80 and base ESM-C along the structure showing the HA1 regions which are improved by fine tuning and H2 regions where performance differs across cleavage states, showing worse performance in the cleaved state.

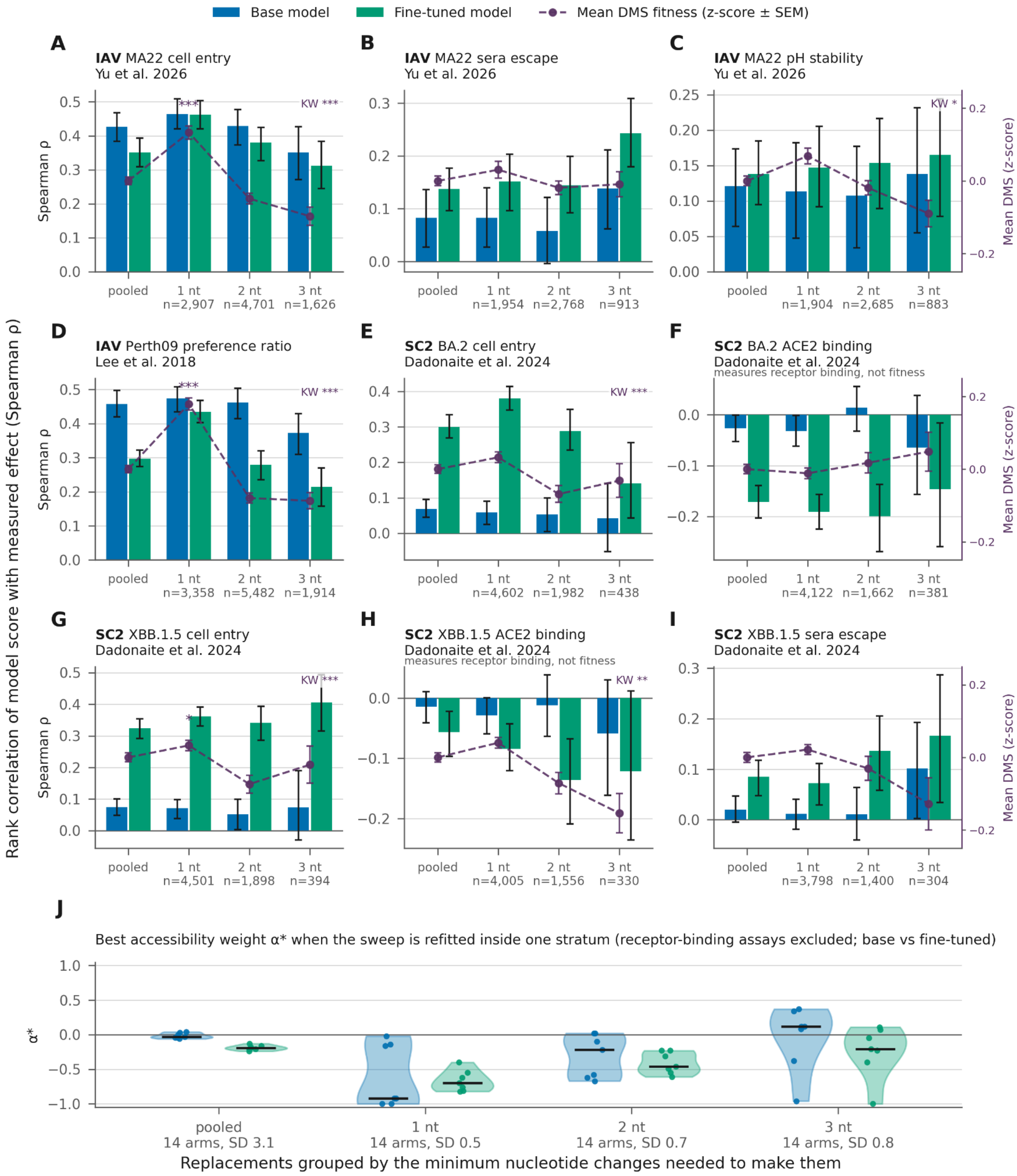


**Figure S7 (A-I)** Spearman ρ between PLM score and DMS rank within codon-distance strata, across Influenza H3N2 and SARS-CoV-2. Overlaid in purple is the mean z-score of the measured phenotype at each codon distance. A Kruskal–Wallis test asks whether the measured effect differs across the three strata: it is significant in six of the nine panels (A, C, D, E, G, H), and the phenotype declines with codon distance in every fitness assay. The four largest effects are influenza cell entry and growth preference (A, D) and SARS-CoV-2 spike-mediated entry (E, G), assays measuring broadly similar phenotypes. Though, pH stability (C) and ACE2 binding (H) are also significant, while the two sera escape assays (B, I) and BA.2 ACE2 binding (F) are not. Accessibility itself carries almost no information about the measured effect once codon distance is held fixed: within strata the median $\rho$ is −0.016 at 1 nt, −0.002 at 2 nt and +0.015 at 3 nt, and no assay reaches $|\rho| = 0.12$ in any stratum. Pooled across strata it is weakly positive (median $\rho$ = +0.019, range −0.004 to +0.108). This is Simpson's paradox — accessibility tracks the phenotype across strata, not within them — and it is what makes the negative $\alpha$ of Figure 4 coherent rather than contradictory: the pooled association is the step count, not accessibility. **(J)** The accessibility weighting $\alpha$ is refitted across and inside each codon distance stratum, see Figure S16 for individual curves. ACE2 assays were not included here due to its inverse shape. Negative $\alpha$ is favoured in 11 out of 14 arms across all replacements and 33 of 42 fitted inside a single stratum. This occurs in 14 of 14 at 1 nt, and 12 of 14 at 2 nt (all 7 fine-tuned models), and only 7 of 14 at 3 nt. This shows down-weighting accessible replacements still helps after codon distance is held fixed. Note that $\alpha$ is not comparable across strata as plotted: fixing the step count removes most of the spread in log(accessibility), so the same real down-weighting requires a larger $\alpha$.

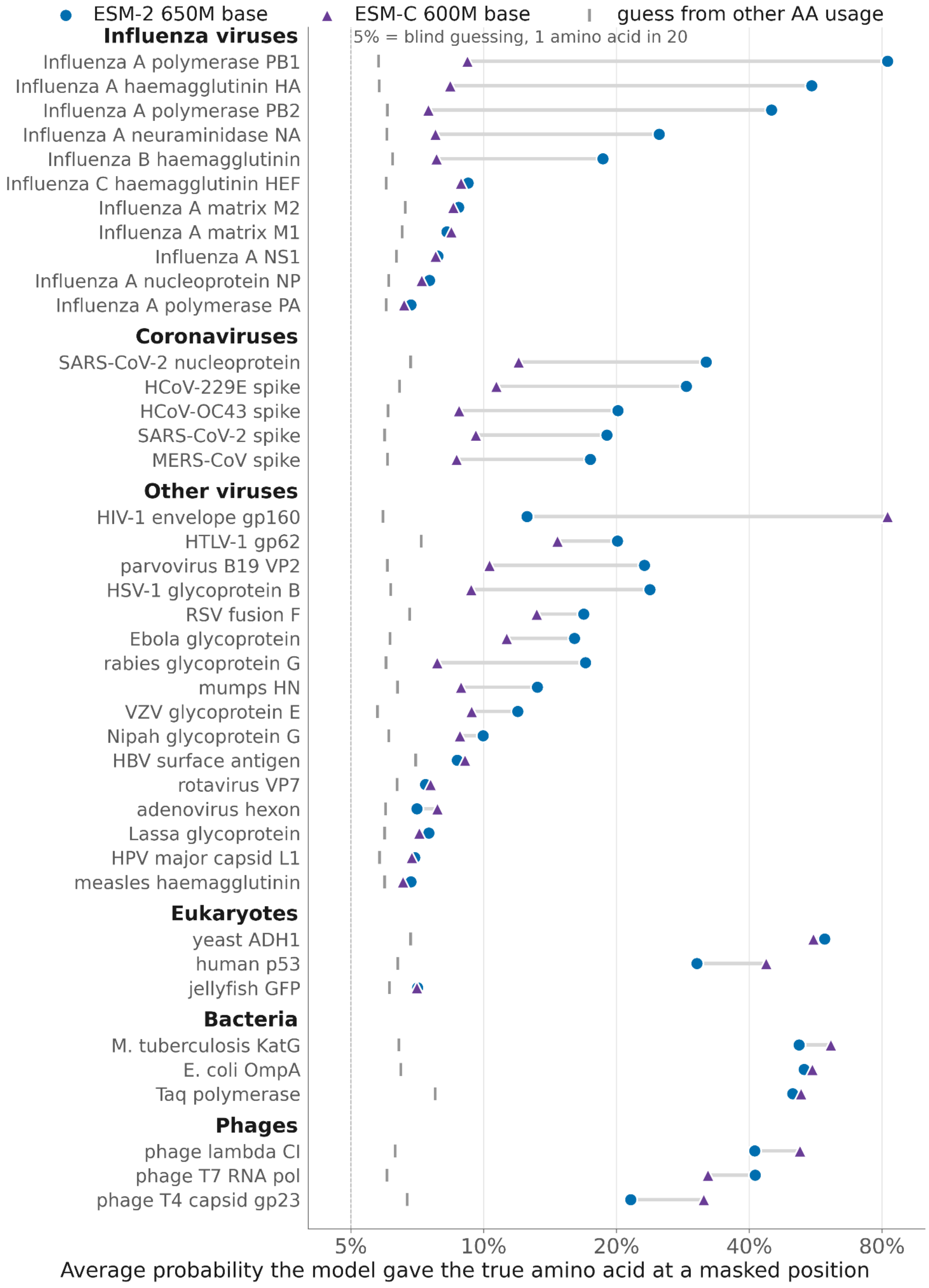

ESM-2 650M base
ESM-C 600M base
guess from other AA usage
5% = blind guessing, 1 amino acid in 20
Influenza viruses
Influenza A polymerase PB1
Influenza A haemagglutinin HA
Influenza A polymerase PB2
Influenza A neuraminidase NA
Influenza B haemagglutinin
Influenza C haemagglutinin HEF
Influenza A matrix M2
Influenza A matrix M1
Influenza A NS1
Influenza A nucleoprotein NP
Influenza A polymerase PA
Coronaviruses
SARS-CoV-2 nucleoprotein
HCoV-229E spike
HCoV-OC43 spike
SARS-CoV-2 spike
MERS-CoV spike
Other viruses
HIV-1 envelope gp160
HTLV-1 gp62
parvovirus B19 VP2
HSV-1 glycoprotein B
RSV fusion F
Ebola glycoprotein
rabies glycoprotein G
mumps HN
VZV glycoprotein E
Nipah glycoprotein G
HBV surface antigen
rotavirus VP7
adenovirus hexon
Lassa glycoprotein
HPV major capsid L1
measles haemagglutinin
Eukaryotes
yeast ADH1
human p53
jellyfish GFP
Bacteria
M. tuberculosis KatG
E. coli OmpA
Taq polymerase
Phages
phage lambda CI
phage T7 RNA pol
phage T4 capsid gp23
5%
10%
20%
40%
80%
Average probability the model gave the true amino acid at a masked position

**Figure S8** ESM-2 and ESM-C's performance on a range of different viral proteins, showing poor performance among many viruses particularly for base ESM-C. We show two nulls, a 5% threshold for random amino acid guesses and a "guess from unmasked amino acid usage" to reflect non-random sequence content and organism specific skews. A few non-viral sequences distributed across other kingdoms are given for some additional context. The jellyfish GFP shows that poor knowledge of PLMs is not fully exclusive to viral sequences.

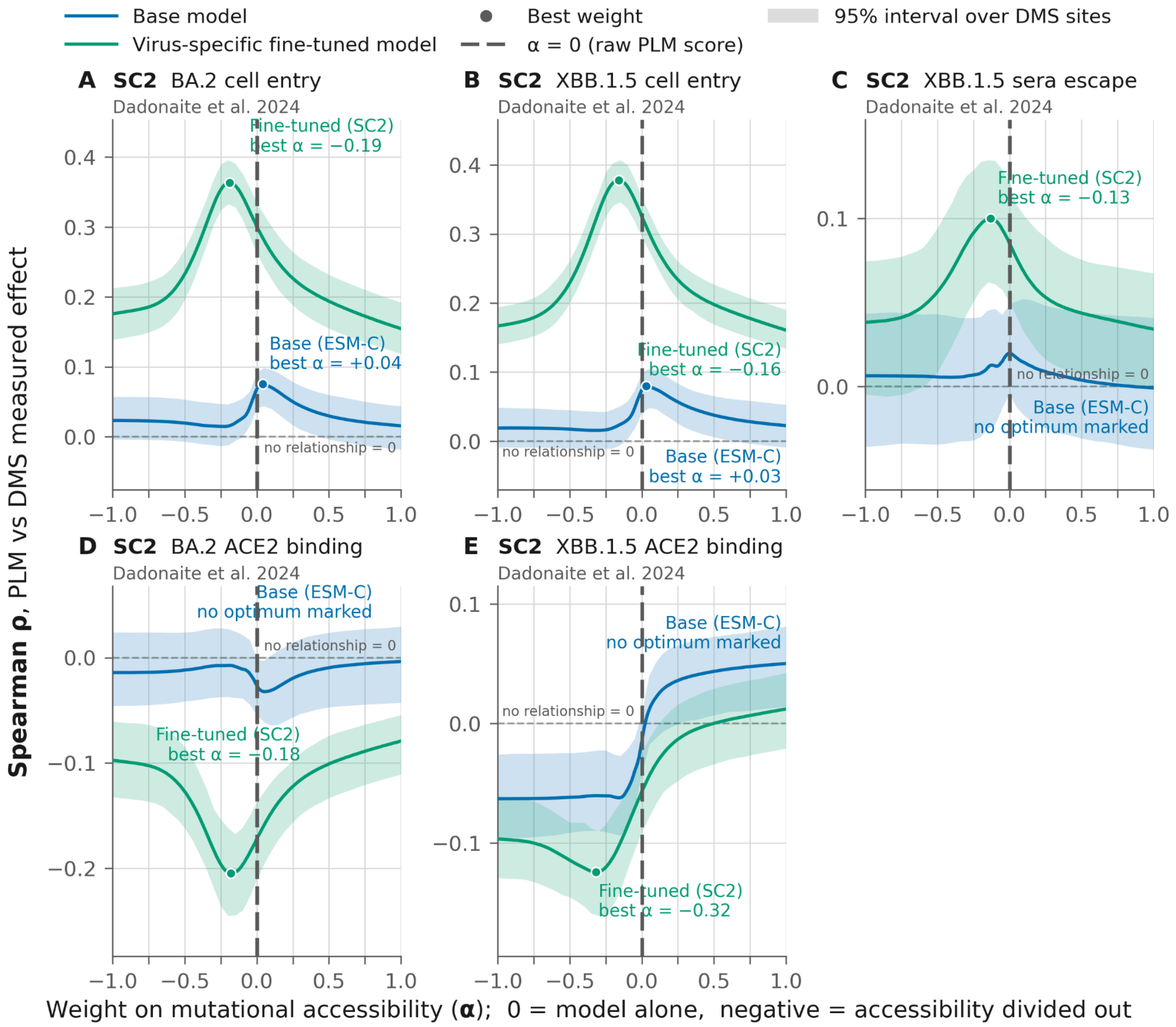


**Figure S9 (A-E)** Deep mutational scanning effect prediction using PLMs for five SARS-CoV-2 datasets across two lineages (Dadonaite et al. 2024). The strongest prediction for the fine-tuned models always comes from a negative α value. Unlike for influenza in Figure 4, the fine-tuned models produce higher absolute Spearman ρ values than the original model for all assays. ESM-C shows very little to no real predictive power of the DMS data, with no benefit of mutational accessibility weighting. This may be due to the poor performance of ESM-C on SARS-CoV-2 spike as seen in Figure S8. The fine-tuned PLM on average gives higher pseudo-probabilities to mutations which reduce ACE2 binding in DMS assays; this may reflect trade-offs with immune escape.

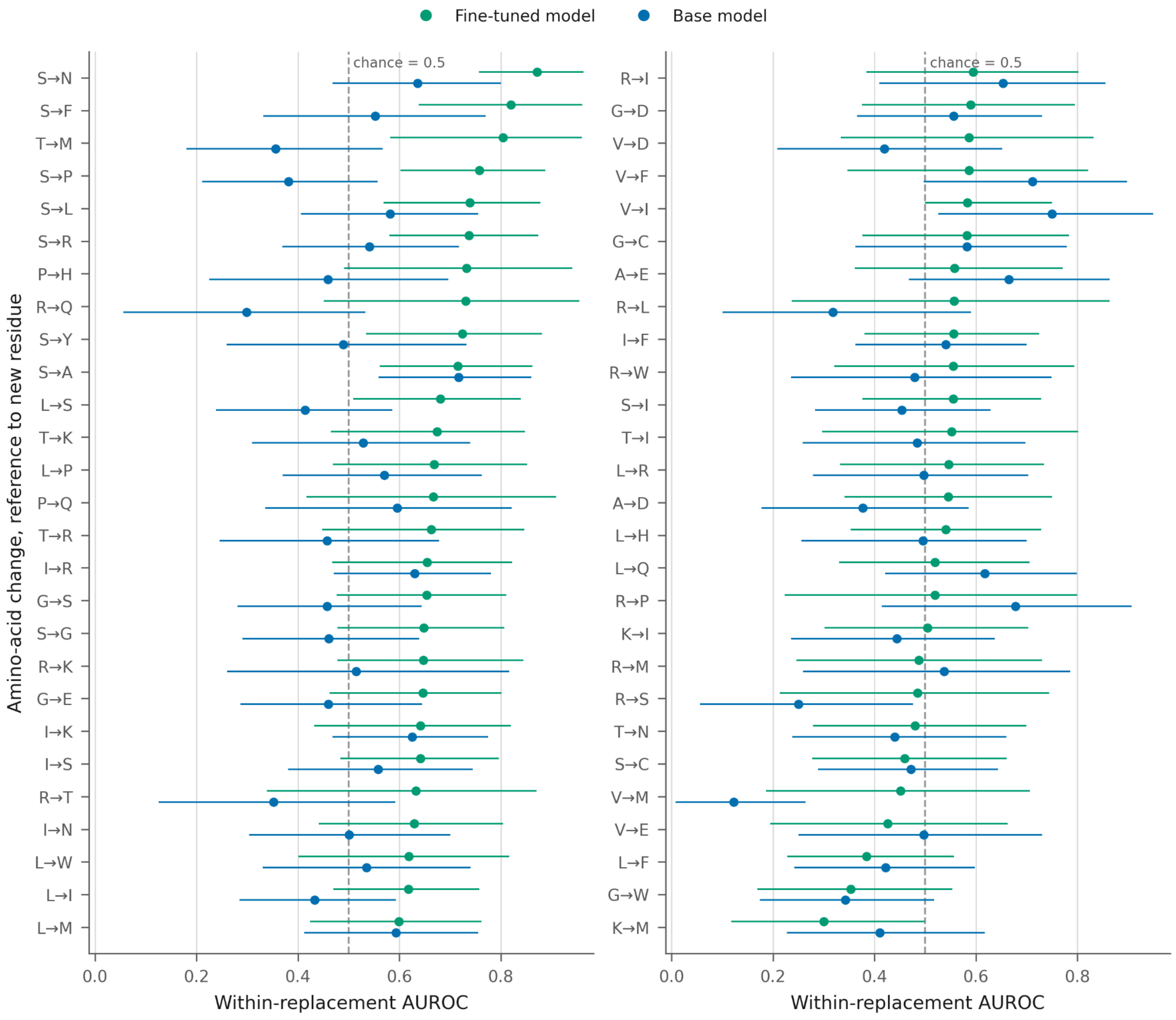


**<u>Figure S10</u>** Asymmetric codon distance amino pairs forest plot showing the PLM learning of H3N2 codon usage (sub-analysis for Figure 5). Within-replacement AUROC for each of the 54 amino-acid changes that have enough sites with one change and with two to three change codons, fine-tuned and base model side by side, strongest fine-tuned first. Whiskers are 95% intervals from resampling sites. 45 of 54 HA-80 fine-tuned values and 25 of 54 base values lie above chance; 9 and 2 amino acid pairs have an interval wholly above 0.5 respectively.

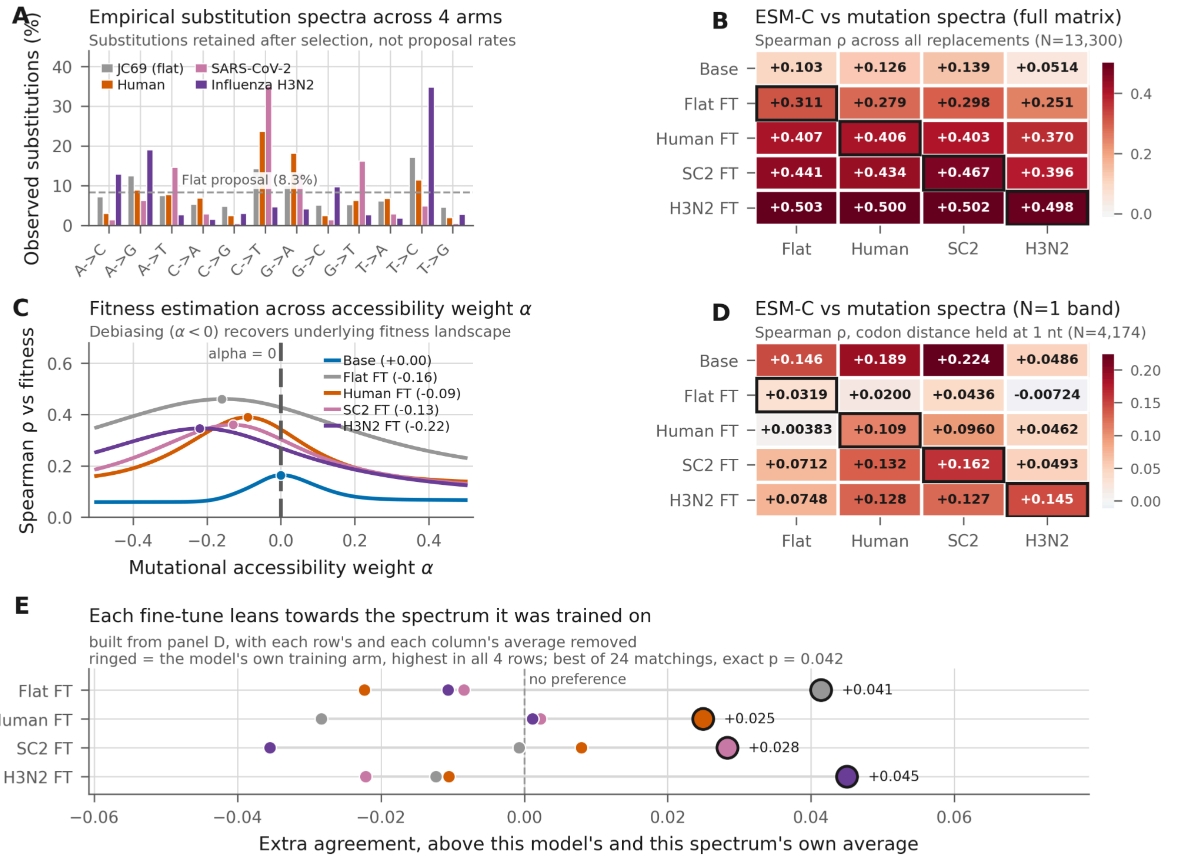


| | Flat | Human | SC2 | H3N2 |
|---|---|---|---|---|
| Base | +0.103 | +0.126 | +0.139 | +0.0514 |
| Flat FT | +0.311 | +0.279 | +0.298 | +0.251 |
| Human FT | +0.407 | +0.406 | +0.403 | +0.370 |
| SC2 FT | +0.441 | +0.434 | +0.467 | +0.396 |
| H3N2 FT | +0.503 | +0.500 | +0.502 | +0.498 |

| | Flat | Human | SC2 | H3N2 |
|---|---|---|---|---|
| Base | +0.146 | +0.189 | +0.224 | +0.0486 |
| Flat FT | +0.0319 | +0.0200 | +0.0436 | -0.00724 |
| Human FT | +0.00383 | +0.109 | +0.0960 | +0.0462 |
| SC2 FT | +0.0712 | +0.132 | +0.162 | +0.0493 |
| H3N2 FT | +0.0748 | +0.128 | +0.127 | +0.145 |

**Figure S11 Fine tuned ESM-C on simulated data (A)** Observed substitution spectra across 12 nucleotide mutation classes in the simulation. Note that JC69 still produces imbalanced observed mutations because of imbalanced starting nucleotide usage and because synonymous mutations are more often transitions than transversions as all twofold sites are separated by transitions, as seen in Xia,Hafner and Sudman (1996). **(B)** correlation of fine-tuned models and the different mutation spectra. **(C)** correlations of fine-tuned models with the simulated fitness landscape. Shows negative α values generate the best correlations with the simulated protein for inference of simulated fitness landscape (fixed across the tree). The best fitness inference is on the neutral mutation spectrum, which is still improved by dividing some mutation accessibility information out, likely because of nucleotide distances. **(D)** N=1 band only, shows the fine-tuned model with neutral mutation spectra correlates least with any mutation matrix **(E)** shows that the models learn their own

N=1 spectra best once the row and column averages are subtracted in panel D

**A**

| | Total | Fewest nucleotide changes needed 1 | 2 | 3 |
|---|---|---|---|---|
| Individual internal branches | 30,028 | 29,875 (99.5%) | 147 (0.5%) | 6 (<0.1%) |
| KY348531 to newest internal nodes | 3,563,404 | 2,930,886 (82.2%) | 537,399 (15.1%) | 95,119 (2.7%) |
| Null, genetic code (J.2; 64 x 20, codon to aa) | 10,754 | 3,334 (31.0%) | 5,486 (51.0%) | 1,934 (18.0%) |
| PLM null, Base model (ESM-2) (J.2, summed probability) | 566 | 364.3 (64.4%) | 174.6 (30.8%) | 27.1 (4.8%) |
| PLM null, HA80 fine-tune (J.2, summed probability) | 566 | 496.3 (87.7%) | 66.4 (11.7%) | 3.3 (0.6%) |

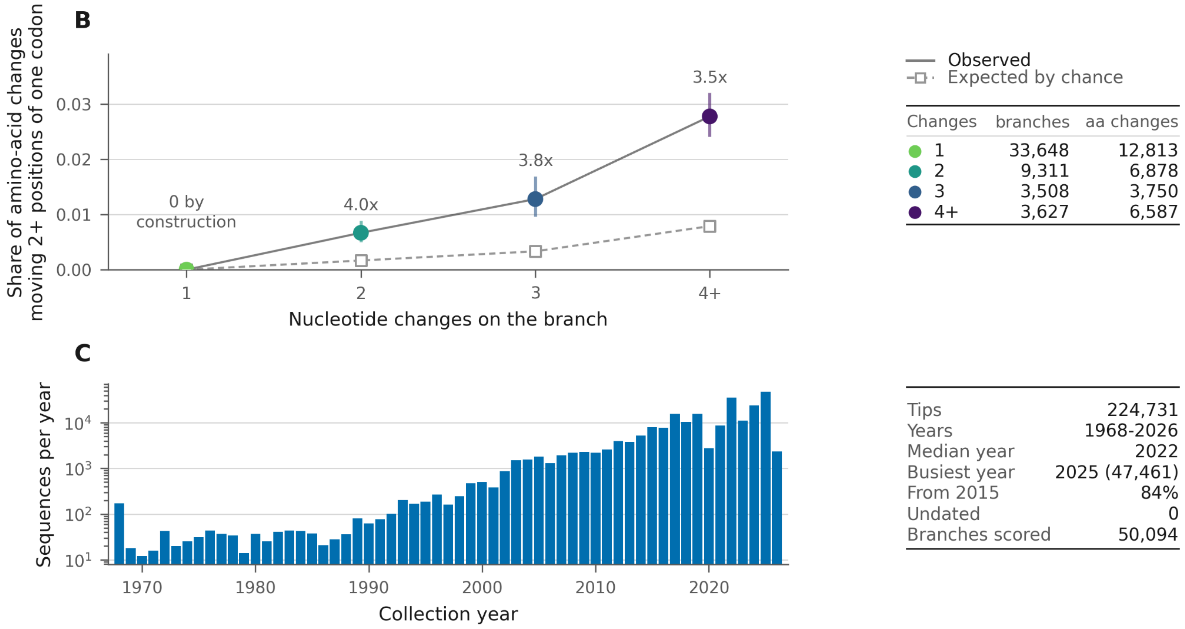


**<u>Figure S12</u>** Analysis of 30,028 individual amino acid replacements across 50,094 internal H2N2 HA branches showing nucleotide mutations **(A)** How many branches contain how many mutations to each codon in the full H3N2 UShER from V-gTK tree expanded from IQ-TREE. Row 1 shows only the internal branches, counting each mutation once in the parsimony tree. Row 2 shows the comparison of an 1968 HA sequence versus all the final internal nodes in the tree to avoid sequencing error singleton tips, which double counts a lot of branch space. Row 3 shows the null expectation from a HA sequence if amino acid replacements were randomly selected from a 20×20 matrix. Rows four and five show the distribution proportional sampling from the PLMs' probabilities would yield.  **(B)** shows the share of amino acid changes which involve more than 1nt change in each branch, with the null below and counts in the legend. **(C)** shows the number of sequences per year in this analysis for context

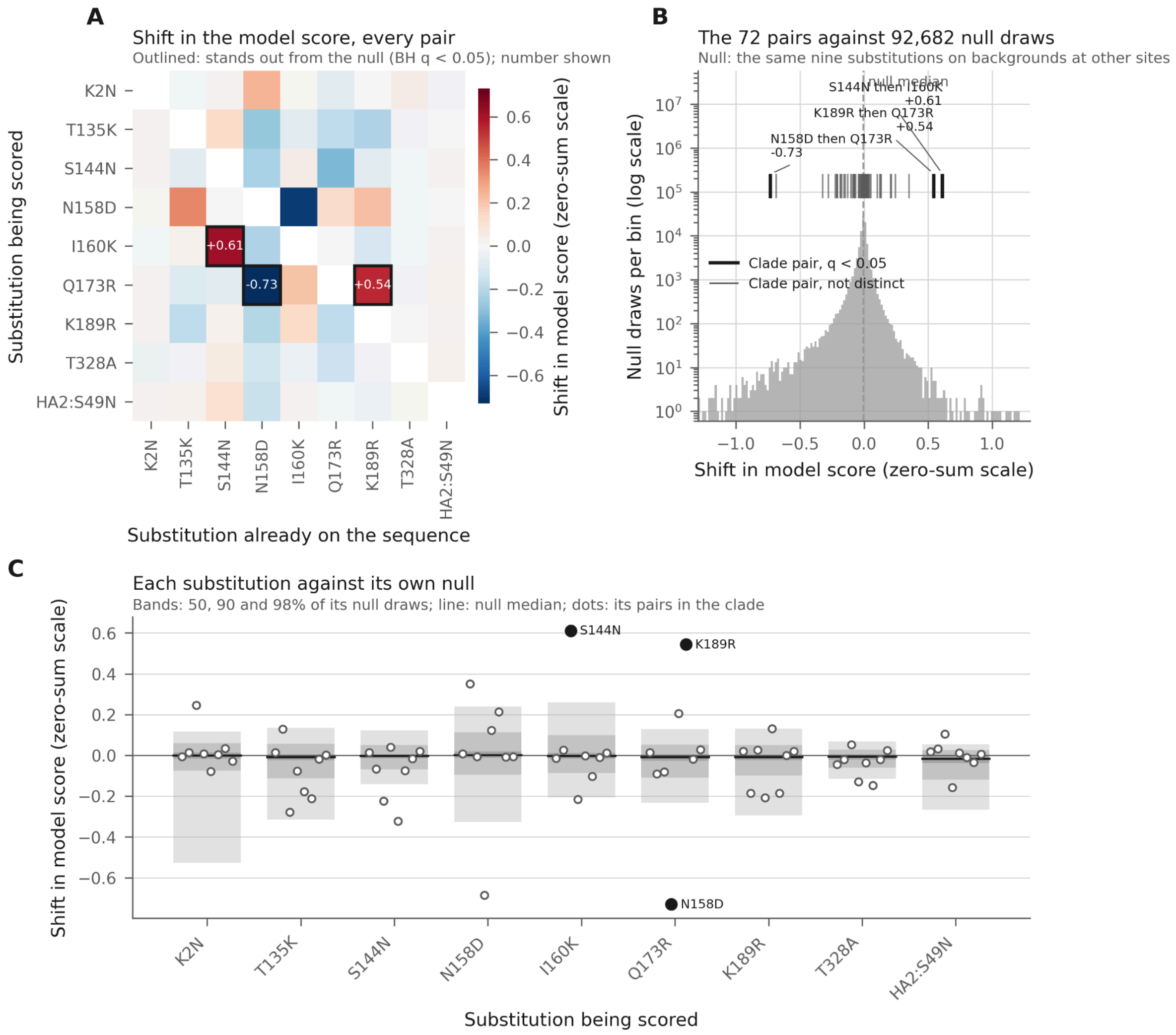


**Figure S13** epistatic scan of K mutations on the J.2 backbone using Zhang zerosum method. It infers both positive and negative epistatic interactions. Some interactions which appear positively epistatic in linear space (increasing in linear probability), sign flip because of changes to the probability of their current amino acid.

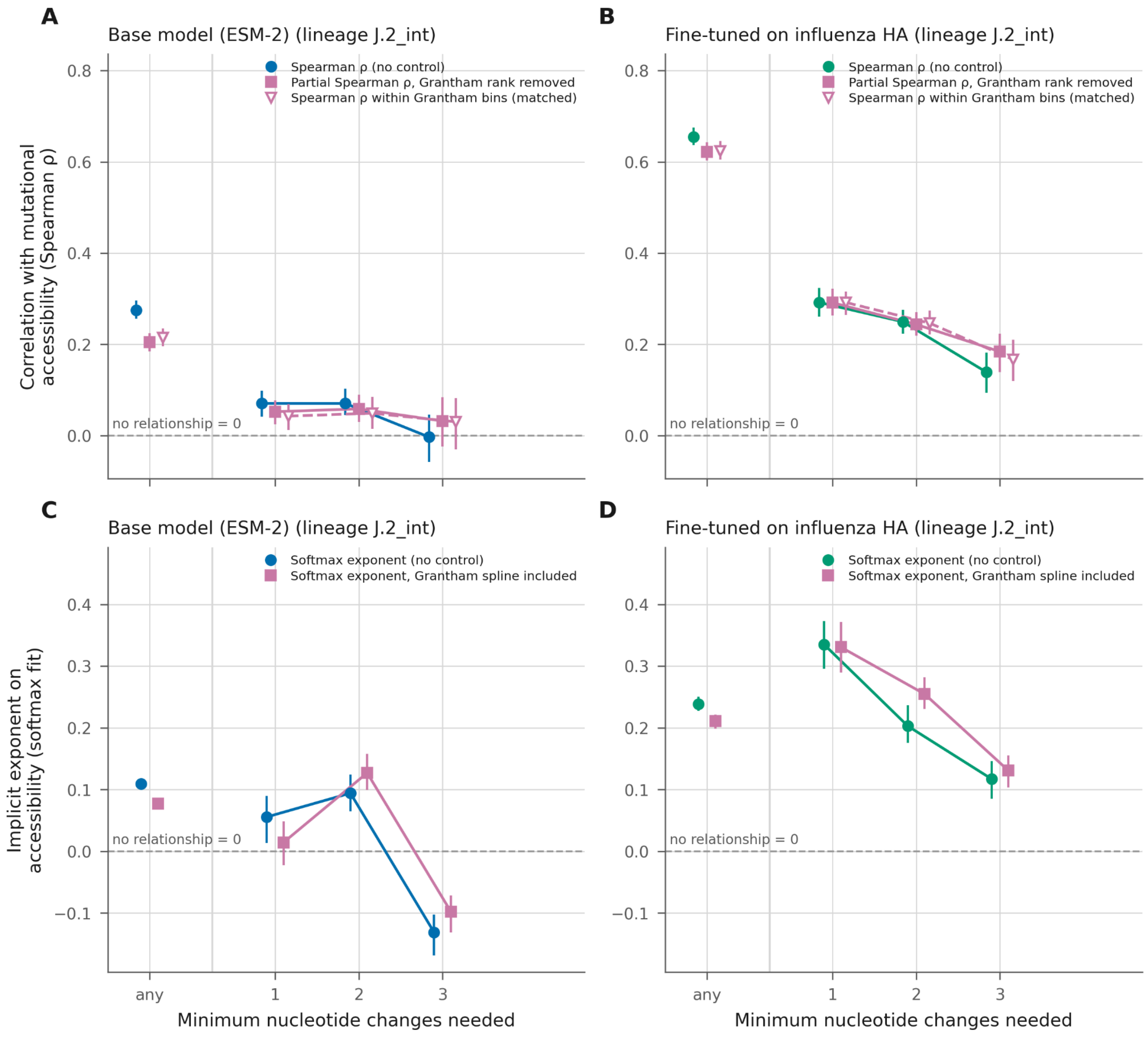


**Figure S14** Alternative methods of adjusting for Grantham's matrix impact on mutational accessibility correlations show biochemical confounding correlations can't explain the PLM correlation with mutational accessibility across the fine-tuned model and across codon distances. **(A and B): rank-based controls.** Partial Spearman rank correlation between ranked model probability ($m$) and ranked mutational accessibility ($a$), controlling for ranked Grantham distance ($c$), for the base model (panel A) and fine-tuned model (panel B). Points show correlations pooled across all possible replacements ('any') and stratified by nucleotide substitution distance (1, 2, or 3 nt changes). To control for non-linear rank associations, a Grantham-matched Spearman is plotted alongside: replacements are partitioned into Grantham distance quintiles, correlations are calculated within each quintile bin, and values are pooled across bins on Fisher's $z$ scale weighted by $n - 3$. In both panels, the 3-nt stratum displays the weakest correlation, turning negative in the base model.

**(C and D)**: likelihood-based controls, in exponent units. Rank correlations weigh every replacement equally: a swap between two variants the model has all but ruled out carries the same weight as a swap between its top two candidates, despite raw model probabilities spanning eight orders of magnitude at a site. Panels C and D eliminate this distortion by modelling the output directly as a within-site softmax:

$$q(site, aa) \propto \exp(\beta \ln a + f(c))$$

where a is mutational accessibility and c is Grantham distance. Chemistry enters as a restricted cubic spline f(c), preventing its effect from being forced into rank-linearity or arbitrary directional smoothness. Fitting within site cancels site-specific intercepts, ensuring estimates depend solely on the relative ordering and log-scale spacing of replacements at that position. Each codon-distance stratum is normalised internally so every site contributes equal weight to its stratum, preventing the 3-change estimate from being dominated by a handful of high-probability sites. The pooled ('any') fit omits codon-distance offsets because discrete steps between distance groups account for the majority of variation in a; including group offsets alongside a common slope would estimate the same quantity twice.
Error bars in all panels indicate 95% confidence intervals derived from 200 bootstraps across sites.

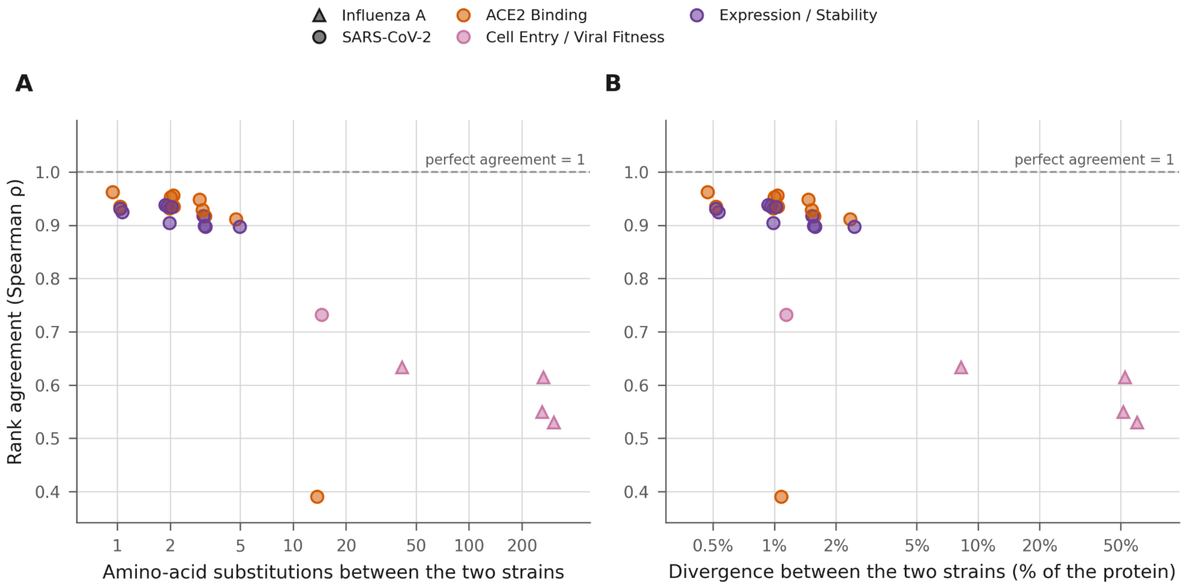


**<u>Figure S15</u>** DMS concordance degrades with protein evolutionary distance, different DMS datasets show correlation decay with divergence.

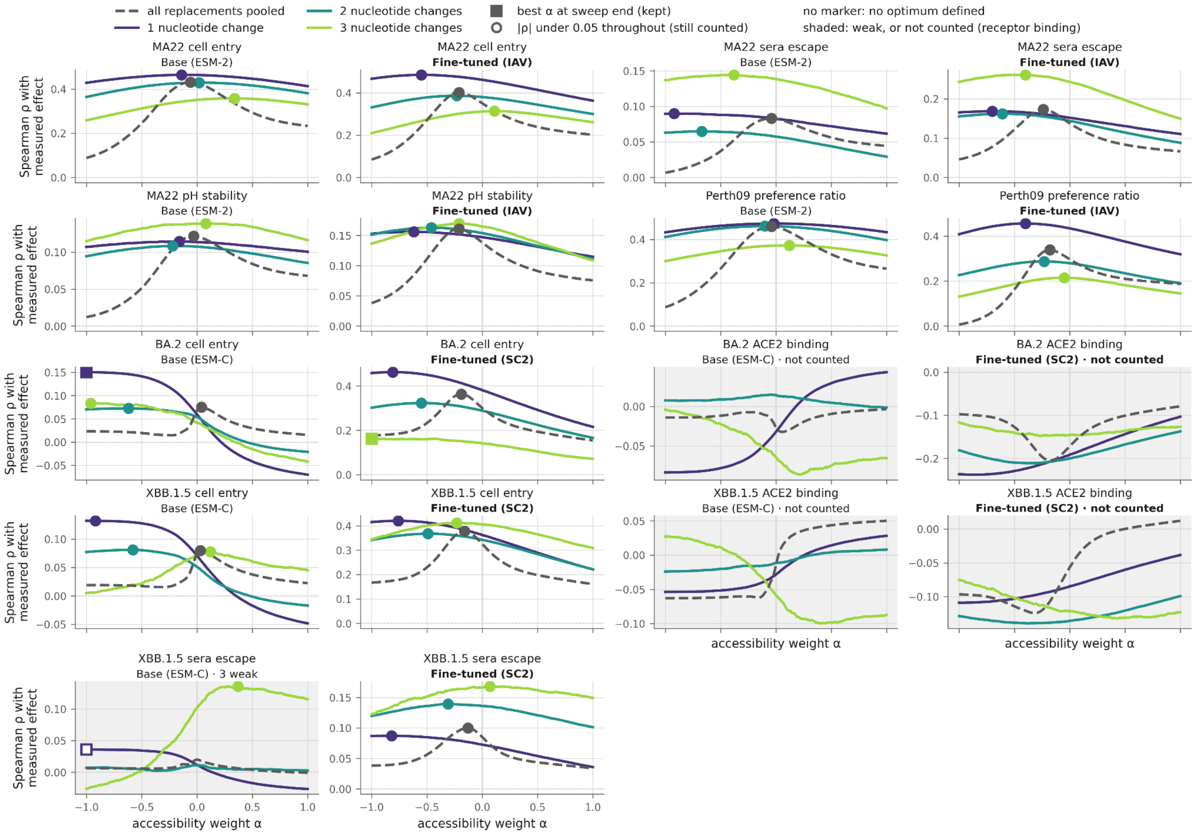


**Figure S16** alpha sweep curves for each individual DMS dataset. Points are shown for peak alpha values, with squares given when the peak is the maximum tested range at -1 or 1 where mutational accessibility is fully divided from or multiplied to the PLM probabilities for correlation with the DMS measured phenotype values. Hollow markers are given when the correlation never reaches 0.05 anywhere in the sweep. The optimum may be perfectly clear, but nothing there is strong enough to lean on. No marker at all was given for the ACE2 dataset curves due to their inverse correlations, and for curves where the weighting moved ρ by less than 0.0005.

The lineages the models are scored on
size, sampling period and amount of variation in each

| Lineage | Sequences | First | Last | Middle 98% | Sites differing | Replacements observed | Homoplasies | Multiple origins | Top 5% freq threshold |
|---|---|---|---|---|---|---|---|---|---|
| C.1 | 5,821 | 2016-05 | 2020-03 | 2017-01 – 2020-02 | 335 | 595 | 251 | 45% | 0.31% |
| E | 1,511 | 2018-01 | 2020-03 | 2018-03 – 2020-01 | 211 | 305 | 87 | 30% | 1.43% |
| D | 1,139 | 2018-11 | 2022-05 | 2019-01 – 2021-09 | 171 | 231 | 65 | 31% | 1.54% |
| E.2 | 1,588 | 2018-09 | 2020-03 | 2019-01 – 2020-03 | 209 | 297 | 87 | 31% | 0.70% |
| G.1 | 1,138 | 2019-11 | 2023-04 | 2021-03 – 2022-12 | 159 | 215 | 47 | 23% | 6.62% |
| G.1.3 | 4,182 | 2020-12 | 2025-02 | 2021-09 – 2023-03 | 247 | 390 | 164 | 45% | 0.49% |
| G.1.1 | 12,597 | 2021-07 | 2024-02 | 2021-10 – 2023-03 | 358 | 704 | 362 | 55% | 0.24% |
| G.2 | 6,969 | 2021-11 | 2024-12 | 2022-03 – 2023-04 | 345 | 609 | 301 | 53% | 0.40% |
| J | 4,059 | 2022-01 | 2025-11 | 2022-06 – 2025-02 | 290 | 488 | 191 | 43% | 0.62% |
| G.2.2 | 2,460 | 2022-07 | 2024-12 | 2022-08 – 2023-03 | 249 | 369 | 131 | 38% | 0.43% |
| J.2 | 26,703 | 2022-12 | 2026-01 | 2023-11 – 2025-10 | 414 | 1,090 | 684 | 66% | 0.30% |
| J.2.4 | 776 | 2024-08 | 2025-12 | 2024-10 – 2025-12 | 135 | 172 | 35 | 22% | 1.26% |
| K | 17,718 | 2025-02 | 2026-02 | 2025-08 – 2026-01 | 399 | 927 | 561 | 64% | 0.29% |
| **All 13** | **86,661** | **2016-05** | **2026-02** | **2017-01 – 2026-01** | **479** | **1,639** | **2,966** | **42%*** | **1.12%*** |

86,661 sequences in all   *  mean of the 13 lineages, not pooled over the sequences

**Supplementary Table 1** The number of sequences from each of the 13 lineages we analysed and the date ranges they were found in. Multiple origins within a lineage were found for an average across lineages of 42% of amino acid replacements, with a higher percentage for lineages with more sequences (Spearman ρ 0.99).

The lineage-defining replacements in the fine-tuning corpus
how much of the new lineage the model had already read

| Replacement | Change | Sequences with it | Share of corpus | First seen | In the human subset | Share of subset |
|---|---|---|---|---|---|---|
| HA1:K2N | K → N | 372 | 5.5% | 1956 | 10 | 0.8% |
| HA1:T135K | T → K | 231 | 2.9% | 1956 | 26 | 2.0% |
| HA1:S144N | S → N | 1,397 | 17.2% | 1957 | 870 | 67.9% |
| HA1:N158D | N → D | 368 | 4.5% | 1942 | 4 | 0.3% |
| HA1:I160K | I → K | 1,764 | 21.8% | 1975 | 1,185 | 92.4% |
| HA1:Q173R | Q → R | 146 | 1.8% | 1902 | 5 | 0.4% |
| HA1:K189R | K → R | 335 | 4.1% | 1977 | 26 | 2.0% |
| HA1:T328A | T → A | 70 | 0.9% | 1974 | 0 | 0.0% |
| HA2:S49N | S → N | 2,345 | 29.0% | 1956 | 1,276 | 99.5% |

8,099 corpus sequences, collected up to 2015-02-27; the subset is human H3N2

**Supplementary Table 2** testing the HA-80 fine-tuning set for the J.2->K mutation set. There are 8,099 sequences, latest collection date 2015-02-27 — all nine of the lineage defining mutations are present. Across all 36 unordered pairs, 33 co-occur; the three that never do are T135K+N158D, T135K+Q173R, Q173R+T328A. No single sequence carries more than five of the nine mutations together.

**Supplementary methods:**

**SC2 fine-tuning**. We used the same implementation as the main paper for re-training the PLM on training data,  MuonW optimiser and MAGMA (Liu et al., 2025; Joo et al., 2026). The training data consists of a panel of diverse coronaviruses alongside SARS-CoV-2 data from 2020-2022.

**UShER and IQ-TREE pan HA tree**

We generated a pan-subtype HA tree using V-gTK, available at https://github.com/centre-for-virus-research/V-gTK (DOI: 10.5281/zenodo.23504636). Query sequences were assigned to their closest segment-4 reference by blastn (BLAST+ 2.17.0; Camacho et al., 2009; -task blastn -max_target_seqs 1), aligned to that reference with Nextalign (2.14.0; Aksamentov et al., 2021), and the resulting per-reference sub-alignments projected onto the coordinate frame of the segment-4 master (AB573800, H1N1) to give a single 572,121-sequence HA alignment.

The backbone was built over MMseqs2 cluster representatives (release 14-7e284; Steinegger & Söding 2017). Clustering was two-step: sequences at ≥90% completeness were clustered with *mmseqs cluster --min-seq-id 0.98 --cov-mode 2 -c 0.8 --cluster-mode 2*, and the remaining partial sequences were then assigned to those clusters by mmseqs easy-search at the same identity threshold, so that no fragment could become a cluster centroid. Greedy incremental clustering (--cluster-mode 2) over a length-sorted input ensures representatives are the most complete member of each cluster. This yielded 4,188 representatives, which together with the 9,470 tips of the previous HA backbone gave a 13,446-sequence backbone alignment.

The backbone phylogeny was inferred with IQ-TREE (v3.0.1; Minh et al., 2020; Wong et al., 2026) under GTR+F+R5 from a parsimony starting tree. The remaining 558,675 sequences were placed onto this starter tree by maximum parsimony using UShER (v0.6.6; Turakhia et al., 2021). Variant calls were generated with faToVcf -includeNoAltN, so that sites uncovered in a partial sequence are encoded as missing rather than silently read as the reference allele. Sequences identical to an existing tip over the aligned region were collapsed onto a single representative before placement. Distinct sequences were placed in batches of 5,000, ordered by informative base count so that the most complete sequences define the structure encountered by later, sparser ones. Placement used usher-sampled with tree optimisation disabled (--optimization_radius 0 --optimization_minutes 0), so samples were added without rearranging existing topology, followed by a repair pass with usher for the small number of tips affected by zero-length-branch collapsing.

The final tree comprises 572,121 tips and 208,626 internal nodes. Of the 18 HA subtypes represented, 15 were recovered as exactly monophyletic clades, with H3 (245,160-tip clade, 0.9992 purity) and H9 (27,491 tips, 0.9999) each containing a handful of non-conforming tips. The tree also includes H18 (n = 9) and H19 (n = 3) which are bat HAs that fall outside the main radiation.

**Simulated fine-tuning.** A synthetic 700-amino-acid founder sequence was evolved under a shared site-wise fitness landscape across four distinct nucleotide substitution regimes: flat JC69 (neutral negative control), human germline, SARS-CoV-2, and influenza A H3N2. ESM-C 600M was scored on the founder background prior to fine-tuning and after fine-tuning on 10,000 evolved sequences from each arm.The synthetic protein (700 aa / 2,100 nt) had no homology to anything real. The founder is not drawn from a real amino-acid composition, it’s sampled from the fitness landscape, so the founder sits at the site optimum at only 66% of sites; median $\pi$ for the founder residue is 0.54. Codons are picked uniformly among synonymous ones, so no codon-usage bias is baked in. Seed 20260924 so the founder is byte-identical across all four arms.

The fitness landscape: 700 × 20 Dirichlet-sampled preference vectors, each row summing to 1, floor 1e-4, this is the ground_truth column in the export. Calibrated to the Perth/2009 HA deep mutational scan from Figure 4 (dms_calibration_targets.json, 566 sites): concentration ~ 0.03·Gamma(k=2.66, θ=0.595)^3.2. Fixation uses proposals from the mutation spectrum and acceptance from the fitness landscape using Halpern–Bruno: $S = \beta(\ln \pi\_new - \ln \pi\_old)$, $\beta = 2.0$, accept with min(1, fix/20). Landscape and β are identical across arms; only the 4×4 nucleotide matrix changes.

Tree shape and distance. A star shaped phylogeny with 10,000 tips evolved independently from the founder. Branch length = accepted amino-acid substitutions / 700 sites, ≈ 0.15 per tip (~106 aa changes, ~2,200–5,400 nt changes, most synonymous). Tips are therefore ~2× that distance from each other, and nothing is shared between them beyond the founder.

Distance was equalised by arm, not by proposal count — star_branch_length (expected proposals/site) is 55 human, 68 SC2, 33 H3N2, 33 JC69, tuned so realised identity to founder lands at 0.87–0.91 in every arm. Mean identity in each simulation arm: human 0.890, SC2 0.907, H3N2 0.906, JC69 0.873.

**Supplementary Section 2**

Supplementary Appendix All genome sequences and associated metadata supporting the findings of this study can be accessed through the persistent digital object identifier https://doi.org/10.55876/gis8.260501cz.

In addition to the minted DOI, GISAID also communicates the aggregation of GISAID accession numbers (EPI_ISL_IDs) through the corresponding EPI_SET_260501cz identifier to facilitate both, the acknowledgment of all data contributors and the direct retrieval of the underlying data from GISAID used in this study.

**Influenza Virus Segments Data Summary**

| Gisaid identifier | DOI | Number of individual viruses | Data Collection range | Number of countries/territories |
|---|---|---|---|---|
| EPI_SET_260501cz | https://doi.org/10.55876/gis8.260501cz | 91,463 | 1905-07-13 to 2026-02-06 | 173 |
